# Systemic low-dose anti-fibrotic treatment attenuates ovarian aging in the mouse

**DOI:** 10.1101/2024.06.21.600035

**Authors:** Farners Amargant, Carol Vieira, Michele T. Pritchard, Francesca E. Duncan

## Abstract

The female reproductive system is one of the first to age in humans, resulting in infertility and endocrine disruptions. The aging ovary assumes a fibro-inflammatory milieu which negatively impacts gamete quantity and quality as well as ovulation. Here we tested whether the systemic delivery of anti-inflammatory (Etanercept) or anti-fibrotic (Pirfenidone) drugs attenuates ovarian aging in mice. We first evaluated the ability of these drugs to decrease the expression of fibro-inflammatory genes in primary ovarian stromal cells. Whereas Etanercept did not block *Tnf* expression in ovarian stromal cells, Pirfenidone significantly reduced *Col1a1* expression. We then tested Pirfenidone *in vivo* where the drug was delivered systemically via mini-osmotic pumps for 6-weeks. Pirfenidone mitigated the age-dependent increase in ovarian fibrosis without impacting overall health parameters. Ovarian function was improved in Pirfenidone-treated mice as evidenced by increased follicle and corpora lutea number, AMH levels, and improved estrous cyclicity. Transcriptomic analysis revealed that Pirfenidone treatment resulted in an upregulation of reproductive function-related genes at 8.5 months and a downregulation of inflammatory genes at 12 months of age. These findings demonstrate that reducing the fibroinflammatory ovarian microenvironment improves ovarian function, thereby supporting modulating the ovarian environment as a therapeutic avenue to extend reproductive longevity.

## Introduction

The ovary is one of the first organs to age in humans. Ovarian follicles, which contain oocytes surrounded by companion somatic cells known as granulosa cells, are the functional unit of the ovary. Primordial follicles are formed before birth and constitute the ovarian reserve or follicular pool which is non-renewable or replenishable and dictates an individual’s reproductive lifespan [1]. Primordial follicles have three fates: remain quiescent, die, or activate and grow [2, 3]. Thus, over time, there is a natural winnowing of ovarian follicle number which is accompanied by a corresponding reduction in gamete quality [4, 5]. Follicles also produce gonadal hormones, such as estrogen and progesterone, which regulate cognitive, cardiovascular, immune, bone, and sexual functions [4]. Thus, ovarian function is critical for overall health. In fact, although women generally have a longer lifespan than men, they spend a longer period of their lives in a frail state due to suboptimal hormone production [6]. Therefore, delaying ovarian aging to extend reproductive longevity and healthspan is currently an unmet clinical need, especially as more women are living longer post-menopause in an altered endocrine environment [7, 8].

In the ovary, follicles develop within a complex microenvironment or stroma enriched in extracellular matrix (ECM) components and heterogeneous cell types, including fibroblasts, endothelial cells, smooth muscle cells, nerve cells, and immune cells among others [9]. With age, the ovary undergoes a shift in immune cell populations characterized by an increase in lymphocytes, including B cells, T cells, and innate-like T cells [10–19]. The aging ovary also assumes a pro-inflammatory state with increased expression and secretion of cytokines such as interleukin-6 (IL6) and tumour necrosis factor alpha (TNFα) [20]. Inflammation typically precedes fibrosis, and ovarian aging is associated with an increase in collagen type I and III and fibrosis-associated proteins as well as a decrease in hyaluronan [20–22]. Many of these age-dependent changes in the stroma are conserved from mouse to humans [21, 23–27]. Finally, increased ovarian fibrosis translates into an organ with quantifiably stiffer biomechanical properties [21].

The age-dependent increase in ovarian fibrosis and stiffness has significant biological consequences, including effects on follicle development and gamete quality as well as ovulation [22, 28, 29]. Moreover, the collagen-rich matrix of the aging ovary may provide a permissive niche for ovarian cancer pathogenesis [23, 30–33]. Thus, targeting ovarian fibrosis is a promising therapeutic intervention to improve ovarian function and extend reproductive longevity. In fact, recent studies have demonstrated that anti-fibrotic drug use in mouse physiologic aging models prevents or reverses ovarian fibrosis [12, 28]. Acute treatment with anti-fibrotic drugs can temporarily restore ovulation, while a long-term treatment of 6 months can modify the ovarian stromal cell population [12, 28]. However, whether low-dose systemic administration of anti-fibrotic and anti-inflammatory drugs beginning prior to advanced reproductive age prevents age-associated fibro-inflammation and extends ovarian function, including follicle number and endocrine activity, has not been systematically investigated.

Therefore, in this study we developed a preclinical pipeline, including *ex vivo* and *in vivo* approaches, to determine whether drugs that target pro-fibrotic molecules and TNF-α, which increase in the ovary with age, can modulate the fibrotic response in ovarian stromal cells and extend parameters of ovarian function. We demonstrate that low-dose systemic treatment of mice with Pirfenidone, an FDA-approved anti-fibrotic drug that is currently used for idiopathic pulmonary fibrosis and reduces TGFβ1 (Transforming Growth Factor Beta 1), PDGF (Platelet-Derived Growth Factor), Collagen type I and type III protein levels in humans, effectively prolonged ovarian function as evidenced by increased follicle numbers, ovulation capacity, gonadal hormone levels, and estrous cyclicity relative to controls [34]. These reproductive longevity phenotypes were associated with a reduction in ovarian fibrosis and inflammation. These pre-clinical data further de-risk the clinical translation of anti-fibrotic agents as a therapeutic intervention to improve ovarian function and reproductive longevity.

## Materials and Methods

### Animals

Female CD1 mice at young (6-12 weeks), mid (6 months), and old (14-17 months) reproductive age were obtained from Envigo (Indianapolis, IN). Mice were housed at Northwestern University’s Center for Comparative Medicine under constant temperature, humidity, and light cycles (12h light/12h night) with access to food and water *ad libitum*. All animal experiments described were approved by the Institutional Animal Care and Use Committee (Northwestern University) and performed in accordance with National Institutes of Health Guidelines.

### Primary ovarian stromal cell isolation and culture

Ovaries from reproductively young mice (N=5 per technical replicate, N=3 replicates per drug) were isolated and placed in dissection media. To enrich for stromal cells and remove most of the oocytes and granulosa cells from large antral follicles, the ovaries were mechanically disrupted with insulin syringes. The remaining stromal enriched tissue was further fragmented into small pieces using forceps and incubated with digestion media comprised of αMEM Glutamax (Thermo Fisher Scientific, Waltham, MA, US) supplemented with 0.5% Pen-Strep, 1% Fetabl Bovine Serum (FBS), 92 digestion units/mL of Collagenase IV (C5138-25MG, Millipore Sigma, Burlington, MA, US) and 0.2 mg/mL of DNAse I (DN25-100MG, Millipore Sigma) at 37°C. The tissue was pipetted every 15 min to facilitate enzymatic digestion. After 30 min, the enzymes in the digestion media were quenched using an equal volume of αMEM Glutamax supplemented with 0.5% Pen-Strep and 10% FBS. The cell suspension was then filtered with a 40 μm strainer to remove undigested tissue, extracellular matrix (ECM), and oocytes, and pelleted at Room Temperature (RT) for 5 min at 1490 x g. The pellet was washed twice with plating media containing RPMI 1640 (Thermo Fisher Scientific) supplemented with 10% FBS and 1% Pen-Strep. The cell number was counted using an Invitrogen automatic counter (Waltham, MA), and a minimum of 0.1×10^6^ cells were plated in 24-well plates with each well containing 750 μl of plating media. Cells were incubated for 24h at 37°C in a humidified atmosphere of 5% CO_2_ in air. After the incubation, cells were treated with: 1) media alone (untreated), 2) 100 ng/ml Lipopolysaccharide (LPS from *Escherichia coli* O55:B5, L2880, Sigma-Aldrich, St. Louis, MO, US), 3) 100 ng/ml LPS and 100 μg/ml Etanercept (Y0002042, Sigma-Aldrich), 4) 100 ng/ml LPS and H_2_O (Etanercept control); 5) 10 ng/ml TGFβ1 (766-MB-005, R&D systems, Minneapolis, MN, US), 6) 10 ng/ml TGFβ1 and 5mM of Pirfenidone (S2907, Selleck Chemicals, Houston, TX, US), 7) 10 ng/ml TGFβ1 and DMSO (Pirfenidone control) for 48h.

### RNA extraction, reverse transcription, and quantitative polymerase chain reaction

Upon treatment, cells were lysed in RLT buffer containing β-mercaptoethanol (RNeasy Micro Kit, Qiagen, Hilden, Germany). RNA was extracted using the same kit, and genomic DNA was digested using on-column DNase I treatment as described previously [35, 36]. RNA quality was assessed using the NanoDrop Microvolume Spectrophotometer (Thermo Fisher Scientific), and the RNA was reserve transcribed to cDNA using the High-Capacity cDNA reverse transcription kit (Thermo Fisher Scientific). Real-Time qPCR (RT-qPCR) was performed to measure *Gusb* (housekeeping gene), or *Tnf, Il6, Il10, Col1a1, and Col3a1* (genes of interest) from control and treated cells using Power SYBR green (Thermo Fisher Scientific). Primers were synthesized by Integrated DNA Technologies (Coralville, IA), and their sequences are in Supplementary Table 1. Primers aligned perfectly to the NCBI Linear mRNA Reference Sequence using the accession numbers found in Supplementary Table 1. Furthermore, nucleotide BLAST confirmed the primers did not have significant identity to other related or unrelated transcripts in mice. Genes of intertest transcripts were normalized to *Gusb* and fold changes were calculated relative to controls (untreated cells) or diluent-treated cells (DMSO or H_2_O) using the 2^−τιτιCt^ method.

### Ovarian explant culture

Ovaries were isolated from mice of mid and old reproductive age (N=3/age), placed in dissection media (L15 (Thermo Fisher Scientific), 0.5% Pen-Strep (Life Technologies, Carlsbad, CA, US), and 1% FBS (Thermo Fisher Scientific) and cut into four even pieces using a scalpel. After washing in growth media (F12/αMEM (Thermo Fisher Scientific) supplemented with 1mg/ml fetuin (Millipore Sigma), 3mg/ml Bovine Serum Albumin (BSA) (Millipore Sigma), 5mg/ml insulin, 5mg/ml transferrin, 5mg/ml selenium (Millipore Sigma) and 10mIU/ml Follicle-stimulating hormone (FSH; obtained from Dr. Mary Zelinski from the Oregon National Primate Research Center (ONPRC)), two ovarian pieces were transferred to the center of a 0.4 μm pore Millicell insert (Millipore Sigma) placed in a 24-well culture plate with each well containing 300 μl of growth media with either 5 mM Pirfenidone, 100 μg/ml f Etanercept, or the respective control (DMSO for Pirfenidone and water for Etanercept). Tissue explants were covered with 4 μl of media and incubated for 24 h at 37°C in a humidified atmosphere of 5% CO_2_ in air. Tissues were imaged in brightfield before and after culture using an EVOS FL Auto Cell Imaging System (Thermo Fisher Scientific). Following the incubation, tissue explants were collected, washed 3 times with Phosphate Buffered Saline (PBS), fixed and processed for histologic analysis as described below.

### Microsurgical placement and removal of osmotic pump

Female CD1 mice were purchased from Envigo at 6 months of age and acclimated in-house for 1 month (N=5 mice per condition and timepoint). The day before of the surgery, 2006 Alzet Osmotic pumps with a mean pumping rate of 0.15 microliters per hour (Alzet, Cupertino, CA) were filled with 10 mg/ml of Pirfenidone or its vehicle (2% DMSO (Millipore Sigma), 30% PEG300 (Selleck Chemicals), and ddH_2_O) and primed in saline solution at 37°C overnight to ensure a constant release of the drugs upon implantation. All these procedures were performed under sterile conditions. Pumps were implanted into mice at 7 months of age using microsurgical techniques at the Microsurgery and Preclinical Research Core (Northwestern University). Mice were anesthetized, and a subcutaneous incision from the dorsal side perpendicular to the spine was made to create a pocket for the pump. Then, an Alzet 2006 pump filled with either Pirfenidone or vehicle was inserted, and the incision was closed using 1-2 surgical clips which were removed 10 to 13 days after surgery. Mice were monitored daily for 15 days. After 6 weeks of treatments, the pumps were removed using the same surgery protocols that were used for pump placement. Following pump removal, mice were aged out for an additional 3.5 months until they were 12 months old. To ensure that all the solution was delivered during the 6 weeks of treatment, the remaining solution of the pump was recovered using the needle provided by Alzet. In all the conditions, less than 10% of the solution remained in the pump.

### Histology, histochemistry, and immunohistochemistry

The left lateral lobe of the liver was isolated from control and Pirfenidone-treated mice, washed three times with warm PBS, and fixed in 10% neutral buffered formalin (Thermo Fisher Scientific) for 24h at RT. Ovaries from control and Pirfenidone-treated mice or ovarian tissue explants were isolated, washed in warm PBS, and fixed with Modified Davidson’s Solution for 3h at RT followed by 4°C overnight (O/N) with gentle rocking. All samples were then washed 3 times in 70% ethanol, processed and paraffin-embedded, and serial-sectioned at a thickness of 5 μm. For hematoxylin and eosin (H&E) and Picrosirius Red (PSR) staining, standard protocols were used [20, 21]. The H&E- and PSR-stained sections were imaged using an EVOS FL Auto Imaging system using a 20X objective and quantifications were performed as described in previous publications [1, 3].

For Cleaved Caspase 3 (CC3; 9579S, Cell Signaling Technology, Danvers, MA, US) detection, tissue sections were deparaffinized in Citrosolv and rehydrated in a series of ethanol dilutions (100, 95, 70, 50%, and a final incubation in distilled H_2_O). Antigen retrieval was performed using 1X Reveal Decloaker (Biocare Medical, Pacheco, CA, US) following manufacturer’s instructions. The slides were then washed twice with Tris-Buffered Saline (TBS) containing 1% TWEEN 20, incubated with 3% hydrogen peroxide (Thermo Fisher Scientific) in TBS and blocked with Avidin and Biotin (Vector Laboratories, Burlingame, CA) for 15 min for each step at room temperature. After a brief rinse in TBS, the slides were blocked in 10% goat serum (Vector Laboratories) and 0.3% Triton-X100 for 1 h. The primary antibody was used at 0.27 μg/mL and incubated O/N at 4°C. Slides were rinsed in TBS-TWEEN 20 three times and incubated in biotinylated rabbit secondary antibody (Vector Laboratories) in TBS-0.3% Triton-X100 for 1.45 h at RT. The slides were washed three times with TBS-TWEEN 20 and incubated 30 min with Avidin-Biotin Complex (ABC) reagent (Vector Laboratories). To amplify the signal, the slides were incubated with TSA Fluorescein System (PerkinElmer, Waltham, MA, US) at 1:50 dilution for 5 min at RT. The slides were mounted using the Vectashield containing DAPI (Vector Laboratories) and imaged using an EVOS FL Auto Cell Imaging System keeping the same settings for all the conditions. As controls, we used non-immune antibodies at the same concentrations. CC3 fluorescence intensity signal per ovarian area was measured, and the fold change was calculated over control conditions. For CC3, three different slides (each one containing three sections) were analyzed from different regions (top, middle and bottom part) of the ovarian tissue explants or ovaries.

### Bone mass density analysis

Mouse bone mass density of control and Pirfenidone-treated mice was measured at the Northwestern University’s Center for Advanced Molecular Imaging (CAMI). Mice were anesthetized and transferred to a dedicated imaging bed in the prone position. Respiratory signals were monitored using a digital monitoring system developed by Mediso (Mediso-USA, Arlington, VA, US). Mice were imaged with a preclinical microPET/CT imaging system (nanoScan PET/CT, Mediso-USA, Arlington, VA, US). CT data were acquired with a 2.2x magnification, <60 µm focal spot, 1 x 4 binning, with 1440 projection views over a full circle, using 70 kVp/88 µA, with a 90 ms exposure time. The projection data were reconstructed with a voxel size of 68 µm^3^ and using filtered (Butterworth filter) backprojection Nucline v3.04 software (Mediso-USA). A bone mineral density (BMD) standard (QRM GmbH, Moehrendorf, Germany) with hydroxyapatite (HA) was used to convert the CT images from Hounsfield units to bone mineral density. The BMD standard was imaged the same day with the same parameters each day.

The reconstructed data were filtered with a non-local means filter and visualized in Amira 2022 (FEI, Houston, TX, US). MRI and microCT images were registered using normalized mutual information. A skeleton region of interest (ROI) from zygomatic arches to one vertebra past the pelvic bone was generated for each mouse by using a 500 HU threshold for larger bones and 300 HU for smaller bones and scapulaes in the CT image. The skeleton was split into upper and lower skeleton at vertebrae L3. The HA standard was segmented with ROIs of 0, 50, 200, 800, and 1200 mg/cm^3^ and used to create a linear correlation between HU and bone density with a r^2^ of 0.99.

### Follicle and corpora lutea (CL) classification and counts

H&E-stained ovarian sections were imaged using an EVOS FL Auto Cell Imaging System. Follicles and CLs were identified morphologically in the ovarian tissue sections as previously performed [22]. The total number of follicles or CLs per total number of sections counted was reported. At least 30 sections encompassing all the ovary were counted.

### Hormone profiling

Estradiol (E2), progesterone (P4) and anti-Müllerian hormone (AMH) were measured from the serum from control and Pirfenidone-treated mice. To obtain serum, 0.5 – 1 mL of blood was first isolated from the superior vena cava and allowed to clot for at least 90 min. The clot was then removed, and the blood centrifuged at 2000 x g at RT for 15 min to isolate the serum. Hormones were measured at the University of Virginia’s Center for Research in Reproduction – Ligand Assay and Analysis Core. E2 levels were measured using the Mouse & Rat Estradiol ALPCO ELISA (#222730) that had a reportable range between 5 – 3200 pg/ml. P4 was measured using the Progesterone Mouse & Rat IBL ELISA (#23K062) with a reportable range between 0.15 and 40 ng/ml. Finally, AMH was measured using an Ansh Labs ELISA with a reportable range of 3.6 to 210.0 ng/ml.

### Estrous cyclicity analysis

Estrous cycle stage was determined in control and Pirfenidone-treated mice by analyzing vaginal cytology and the morphology and proportion of the different cell types, as previously described [37]. Vaginal cells were obtained daily between 10 am and 11 am for a total of 15 days through gentle lavage of the vagina using 0.9% sterile saline solution. The cells were then imaged by transmitted light microscopy using an EVOS FL Auto Cell Imaging System (Thermo Fisher Scientific). Animals that were able to transition through all 4 estrous cycle stages (proestrus, estrus, metestrus, and diestrus) during the 15 days of analysis were classified as “cycling,” and those that did not complete a full estrous cycle during 15 days were classified as “non-cycling.”

### Bulk ovary RNA-sequencing and bioinformatics analysis

The left ovaries of control and Pirfenidone-treated mice were isolated, washed with warm PBS, and incubated in RNAlater Stabilizing Solution (Thermo Fisher Scientific) O/N at 4°C. The RNAlater was then removed, and the ovaries were snap-frozen on dry ice and stored at −80°C. To lyse the ovaries, samples were thawed on ice and lysed in RLT buffer with β-mercaptoethanol (RNeasy Micro Kit, Qiagen) using a bead homogenization system (Thermo Fisher Scientific). RNA was then isolated using the RNeasy Micro Kit as previously described. The extracted RNA was submitted to the Northwestern University NUseq Core Facility to conduct the stranded total RNA-seq. Briefly, total RNA samples were checked for quality on the Agilent Bioanalyzer 2100, and quantity was assessed with the Qubit fluorometer. The Illumina Stranded Total RNA Prep Kit was used to prepare sequencing libraries; the procedure was performed without modifications. This procedure includes rRNA depletion, remaining RNA purification and fragmentation, cDNA synthesis, 3’ end adenylation, Illumina adapter ligation, library PCR amplification, and validation. An lllumina HiSeq 4000 Sequencer was used to sequence the libraries with the production of single-end, 50 bp reads.

The quality of DNA reads, in fastq format, was evaluated using FastQC. Adapters were trimmed, and reads of poor quality or aligning to rRNA sequences were filtered. The cleaned reads were aligned to the *Mus musculus* genome (mm10) using STAR [38]. Read counts for each gene were calculated using htseq-count [39] in conjunction with a gene annotation file for mm10 obtained from UCSC (University of California Santa Cruz; http://genome.ucsc.edu). Normalization and differential expression were determined using DESeq2 [40]. A false discovery rate (FDR)-adjusted p-value less than 0.05 was used as the cutoff for determining significantly differentially expressed genes. Pathway analysis was performed on both gene lists using GeneCoDis [41–43] and ShinnyGO 0.80 [44] to identify pathways enriched in upregulated and downregulated genes.

### Measurement of systemic cytokines, chemokines, and growth factors

We measured 48 cytokines, chemokines, and growth factors in the serum of control and Pirfenidone-treated mice with a Luminex multiplex technique using the ProcartalPlex^TM^ Mouse Immune Monitoring Panel, 48plex (EPX480-20834-901, Thermo Fisher Scientific) according to the manufacturer’s instructions. Undiluted serum samples were analyzed by the Northwestern University’s Comprehensive Metabolic Core.

### Statistics

The normality of the samples was assessed using the Shapiro-Wilk test. Normally distributed samples were analyzed using Student t-test or one-way ANOVA for comparisons between 2 groups or more than 2 groups, respectively. For non-normally distributed samples, we used the non-parametric Mann-Whitney test. Variability within the experimental groups is reported as standard error of mean (SEM). All analyses were performed using the statistical package in Prism 9.4 software (GraphPad Software).

## Results

### Pirfenidone but not Etanercept can modulate fibroinflammatory gene expression in primary ovarian stromal cells

To identify drugs that can modulate the expression of inflammatory and/or fibrotic genes in the ovary, we developed an *ex vivo* screening pipeline using primary ovarian stromal cells. Given that TNFα expression and secretion increase significantly in the ovary with age and is a central mediator of pro-inflammatory responses, we first tested the efficacy of Etanercept, an FDA-approved TNF inhibitor used for the treatment of autoimmune conditions such as rheumatoid arthritis that binds to TNFα and TNFβ and prevents their activation of the inflammatory cascade [45, 46]. Primary ovarian stromal cells from reproductively young mice were isolated and cultured in the presence of LPS for 24h to induce an inflammatory response as has been reported previously in other cells [47–50] (Figure 1A). LPS induced a pro-inflammatory response as evidenced by increased expression of *Tnf* (p≤0.01; Figure 1B). However, co-treatment of LPS-induced cells with Etanercept for 48h did not decrease the expression of *Tnfα* or another inflammatory gene, *Il6*, or the anti-inflammatory gene *Il10*, relative to controls treated with diluent alone (Figure 1A and B). These findings demonstrate that Etanercept does not prevent LPS-induced inflammation in primary ovarian stromal cells under the conditions evaluated.

**Figure 1:**
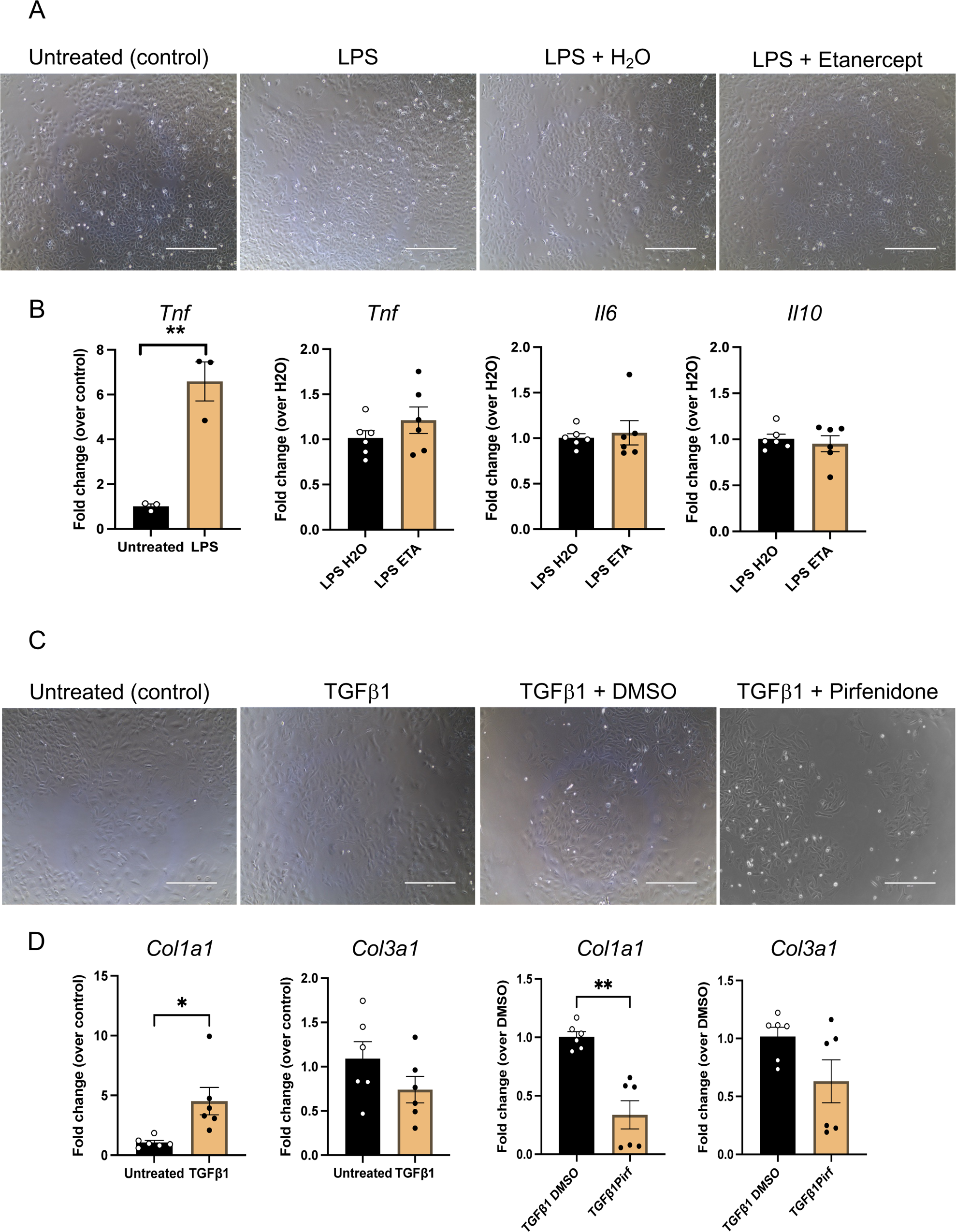
Etanercept does not impact the expression of inflammatory markers, whereas Pirfenidone reduces *Col1a1* expression in primary ovarian stromal cells. (A) Representative transmitted light images of primary ovarian stromal cells that were either untreated or treated with LPS, LPS and H_2_O, or LPS and Etanercept (ETA) for 48 h. (B) Quantification of the relative expression of *Tnf*, *Il6,* and *Il10* transcripts in untreated cells or those treated with LPS, LPS and H_2_O, or LPS and Etanercept. (C) Representative transmitted light images of primary ovarian stromal cells that were either untreated or treated with TGFβ1, TGFβ1 and DMSO, or TGFβ1 and Pirfenidone for 48 h. (D) Quantification of the relative expression of *Col1a1* and *Col3a1* transcripts in untreated cells or those treated with TGFβ1, TGFβ1 and DMSO, or TGFβ1 and Pirfendione. N=3 independent stromal cell cultures per condition. Significant differences are noted (* indicates p<0.05 and ** indicates p<0.01). The scale bar is 400 μm.

Given that fibrosis, caused by the excess accumulation of collagen I and III, is one of the hallmarks of ovarian aging, we tested whether an anti-fibrotic drug, Pirfenidone, could inhibit the fibrotic response in primary ovarian stromal cells [20]. Pirfenidone is an FDA-approved, broad-spectrum anti-fibrotic drug used in the treatment of pulmonary idiopathic fibrosis and functions by decreasing Type I and Type III collagen, PDGF, and TNFα protein levels [51–53], and the expression of these molecules increases with age in the ovary [20, 21]. TGFβ is a pro-fibrotic stimulus and induces collagen expression in various cell types *in vitro* such as cardiac fibroblasts, alveolar fibroblasts, and myoblasts [54]. In primary ovarian stromal cells, TGFβ1 induced *Col1a1* but not *Col3a1* transcripts (p≤0.05, Figure 1 C and D). Co-treatment of TGFβ1-stimulated primary ovarian stromal cells with Pirfenidone decreased *Col1a1* but not *Col3a1* expression relative to controls treated with diluent alone (p≤0.01, Figure 1 C and D). Given the ability of Pirfenidone to abrogate the pro-fibrotic response of primary ovarian stromal cells *ex vivo,* we wanted to further confirm the safety profile of this drug on ovarian tissue prior to performing *in vivo* studies. Therefore, we cultured ovarian tissue explants from ovaries isolated from mice of mid- and old-reproductive age in the presence or absence of Pirfenidone. Pirfenidone treatment neither altered tissue explant architecture nor increased apoptosis relative to controls cultured in vehicle alone (Supplementary Figure 1). These data demonstrate that Pirfenidone attenuates *Col1a1* expression induced by a pro-fibrotic stimulus, and it does not exhibit overt signs of toxicity in the ovary. These findings provided a strong rationale for advancing Pirfenidone into *in vivo* preclinical studies.

### Osmotic pumps can be used to deliver Pirfenidone systemically to study its effect on ovarian aging

Fibrosis and inflammation are first evident in CD1 mice at 7 months of age [20]. Therefore, we sought to develop a preclinical *in vivo* model to determine whether systemic delivery of Pirfenidone at this timepoint could delay age-related fibro-inflammation and improve markers of ovarian function (Figure 2A). At 7 months of age, osmotic pumps filled with either Pirfenidone or vehicle were implanted subcutaneously in mice, and the drug was delivered systemically at a constant flow rate for 6 weeks (Figures 2A-C). We validated that the pump was able to effectively achieve systemic delivery by implanting pumps filled with 1% Evans Blue, a non-toxic dye that can easily be visualized by its blue color (Figure 2B). By 3 days post-pump placement, the blue dye was observed underneath the skin throughout the body (Figure 2B), and after a week, the dye was observed in the mouse limbs.

**Figure 2:**
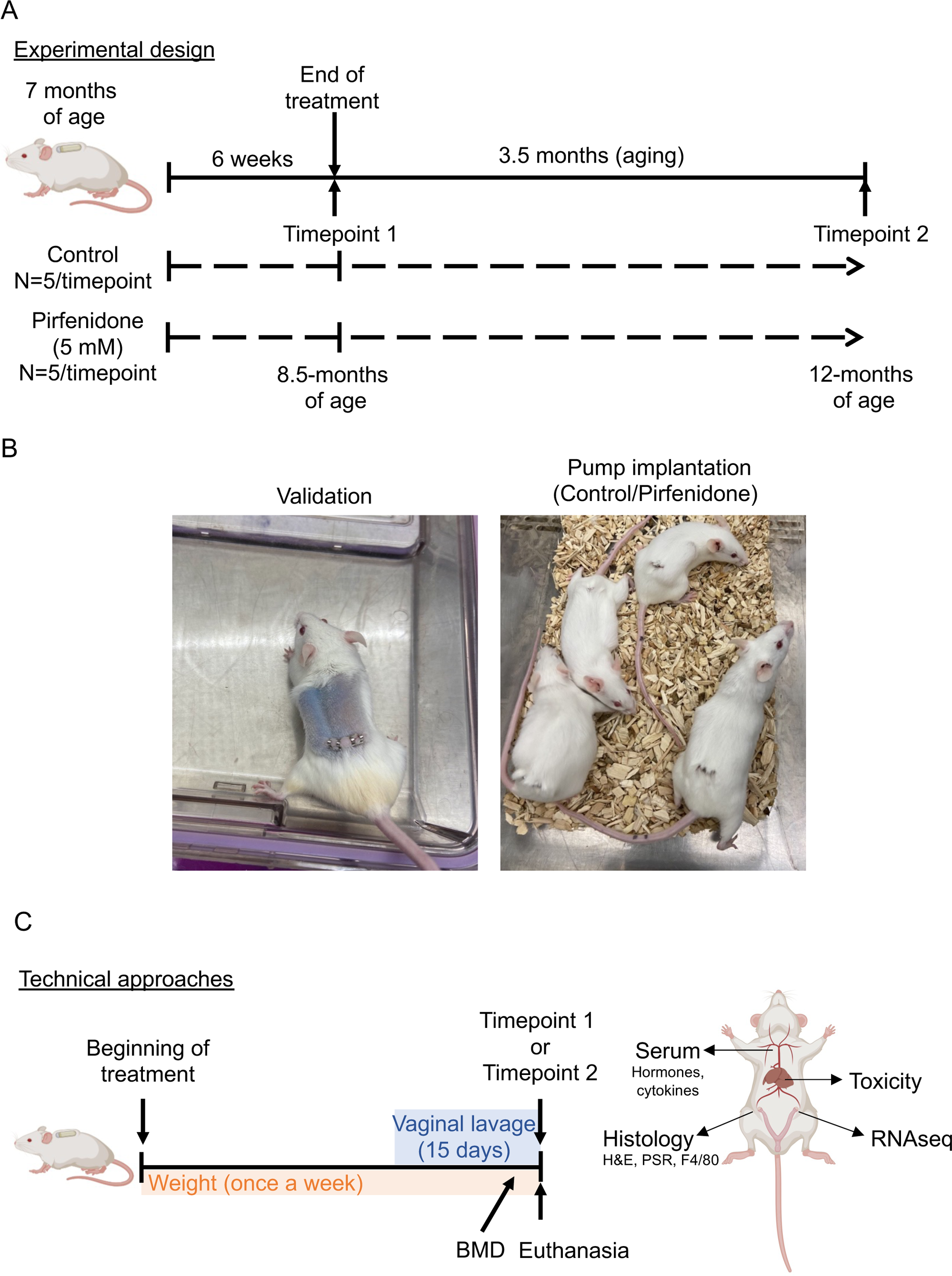
Paradigm of *in vivo* drug delivery and experimental reproductive and general health endpoints. (A) Mini-osmotic pumps filled with diluent (30% PEG, 3% DMSO and H_2_O) or 10 mg/ml Pirfenidone were implanted in CD1 female mice at 7 months of age, and systemic delivery was performed for 6 weeks. At the end of the treatment (Timepoint 1), one cohort of mice was analyzed (8.5 months), whereas another cohort was aged for an additional 3.5 months following removal of the mini-osmotic pump. This cohort of mice was then analyzed at 12 months (Timepoint 2). (B) The left image shows a 7-month old CD1 female mouse bearing a mini-osmotic pump containing non-toxic Evans blue dye. The distribution throughout the body demonstrates effective systemic delivery. The right image shows 7-month old CD1 mice implanted with mini-osmotic pumps that release either control solution (vehicle) or Pirfenidone (10 mg/ml). (C) In each experimental condition, mice were weighed weekly. Fifteen days prior to Timepoint 1 or Timepoint 2, estrous cyclicity was monitored by daily vaginal lavage. The week before Timepoint 1 or Timepoint 2, microCT scans were performed to assess bone mass density (BMD). At euthanasia (Timepoint 1 and Timepoint 2), blood was extracted to isolate serum for hormone and cytokine analyses, the left ovary was isolated for RNA extraction and RNAseq analysis, the right ovary was isolated and fixed for histological and immunohistochemical analyses, and the liver was isolated and fixed for toxicity analysis.

Following the Pirfenidone treatment, one cohort of mice was analyzed immediately (Timepoint 1, 8.5-months of age), whereas a second cohort was aged an additional 3.5 months post-treatment after pump removal (Timepoint 2, 12-months of age) (Figure 2A). A series of assays were then performed to assess parameters of overall and reproductive health. Mouse weight was tracked weekly across the study, and fifteen days prior Timepoint 1 or 2, estrous cyclicity was monitored daily by assessing vaginal cytology. Immediately before Timepoint 1 and 2, bone mass density measurements were performed using microCT scanning. At the time of euthanasia, blood was collected for hormone and cytokine measurements, the right ovary and liver from each mouse was processed for histologic analysis, and the left ovary from each mouse was isolated for bulk RNAseq analysis (Figure 2C).

### Pirfenidone treatment mitigated ovarian fibrosis without affecting overall health parameters

To validate whether our Pirfenidone treatment paradigm modulated the collagen content of mouse ovaries, we performed PSR staining (Figure 3A). As expected, there was an age-dependent increase in collagen in control mice, with a 13.3-fold increase between 8.5 months and 12 months of age (Figure 3B-C). Although Pirfenidone-treated mice also exhibited an age-dependent increase in ovarian collagen, the magnitude was less than that observed in controls, with only a 3.6-fold increase occurring between 8.5 months and 12 months of age (p≤0.01, Figure 3B-C). Thus, Pirfenidone treatment mitigates the age-dependent increase in ovarian fibrosis.

**Figure 3:**
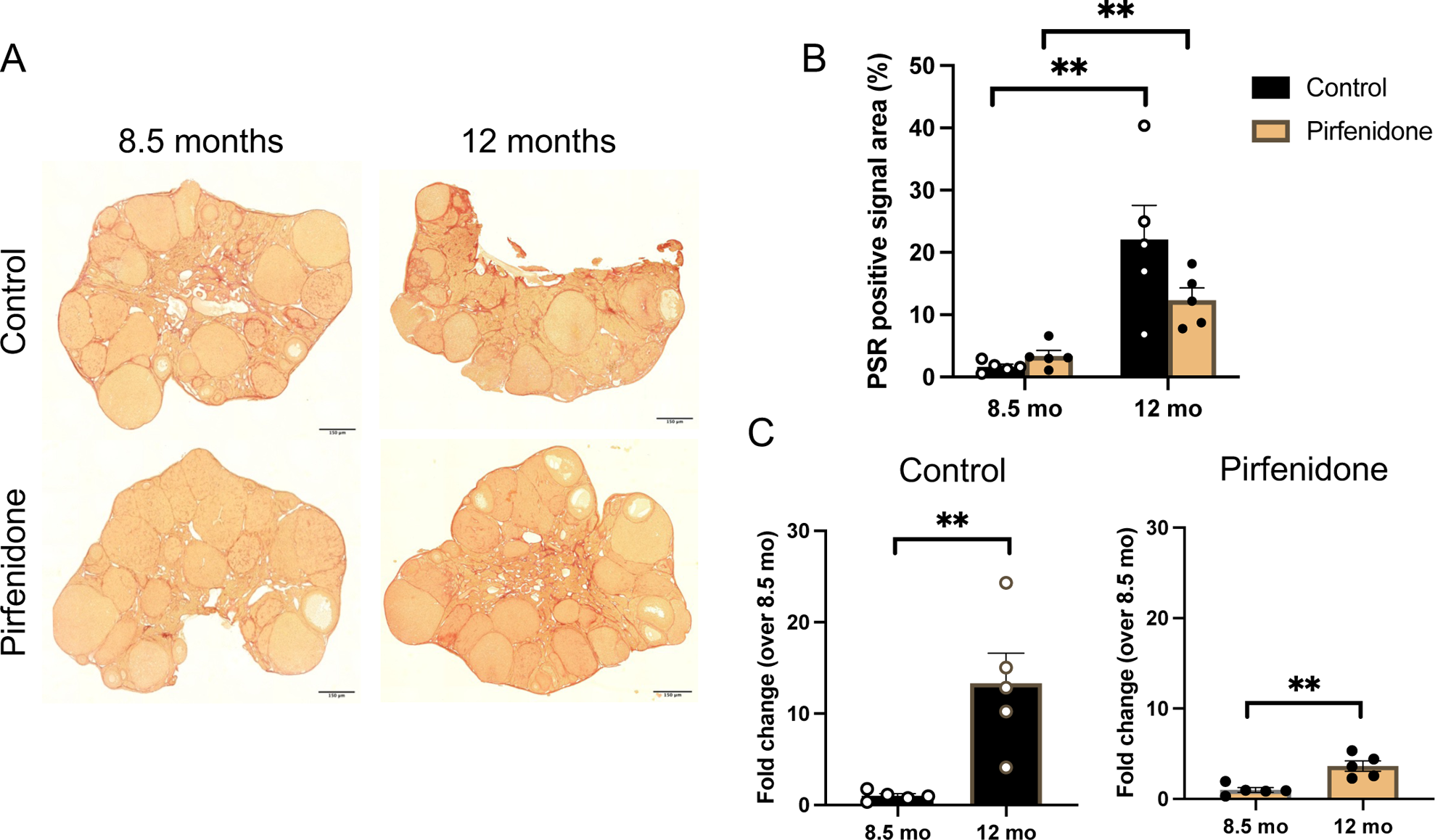
Pirfenidone-treated mice exhibit an attenuation in the age-associated increase in ovarian collagen relative to controls. (A) Representative histological sections of ovarian tissue stained with PSR from control mice or those treated with Pirfenidone at 8.5 months and 12 months. The scale bar is 150 μm. (B) The quantification of the percentage of PSR-positive area per section is shown in control and Pirfenidone-treated mice at 8.5 and 12 months. (C) Graphs represent the fold change of collagen content at 12 months over 8.5 months in control (left) and Pirfenidone-treated mice (right). Significant differences are noted (** indicates p<0.01).

To examine whether there was any overt toxicity or general health effects of systemic Pirfenidone treatment, we assessed body weight, liver histology, and bone mass density. During the 6-week treatment period, there were no differences in body weight between experimental cohorts (Figure 4A). After treatment, mice in the control group maintained a fairly constant body weight throughout the experiment, with a few mice exhibiting a reduction – a phenomenon commonly observed in aged mice [55]. However, those treated with Pirfenidone exhibited a steady increase in weight, reaching a significant difference at weeks 15 and 16 (p≤0.05), although no significant differences were detected at 12 months of age (Figure 4B). Since Pirfenidone is associated with liver injury in humans [56, 57], we performed a histologic evaluation of liver tissue from both control and Pirfenidone-treated mice (Supplementary Figure 2). We did not observe morphologic differences with respect to liver cellular architecture, steatosis, necrosis, or inflammatory infiltrate, indicating that the administered treatment paradigm of Pirfenidone did not induce liver damage.

**Figure 4:**
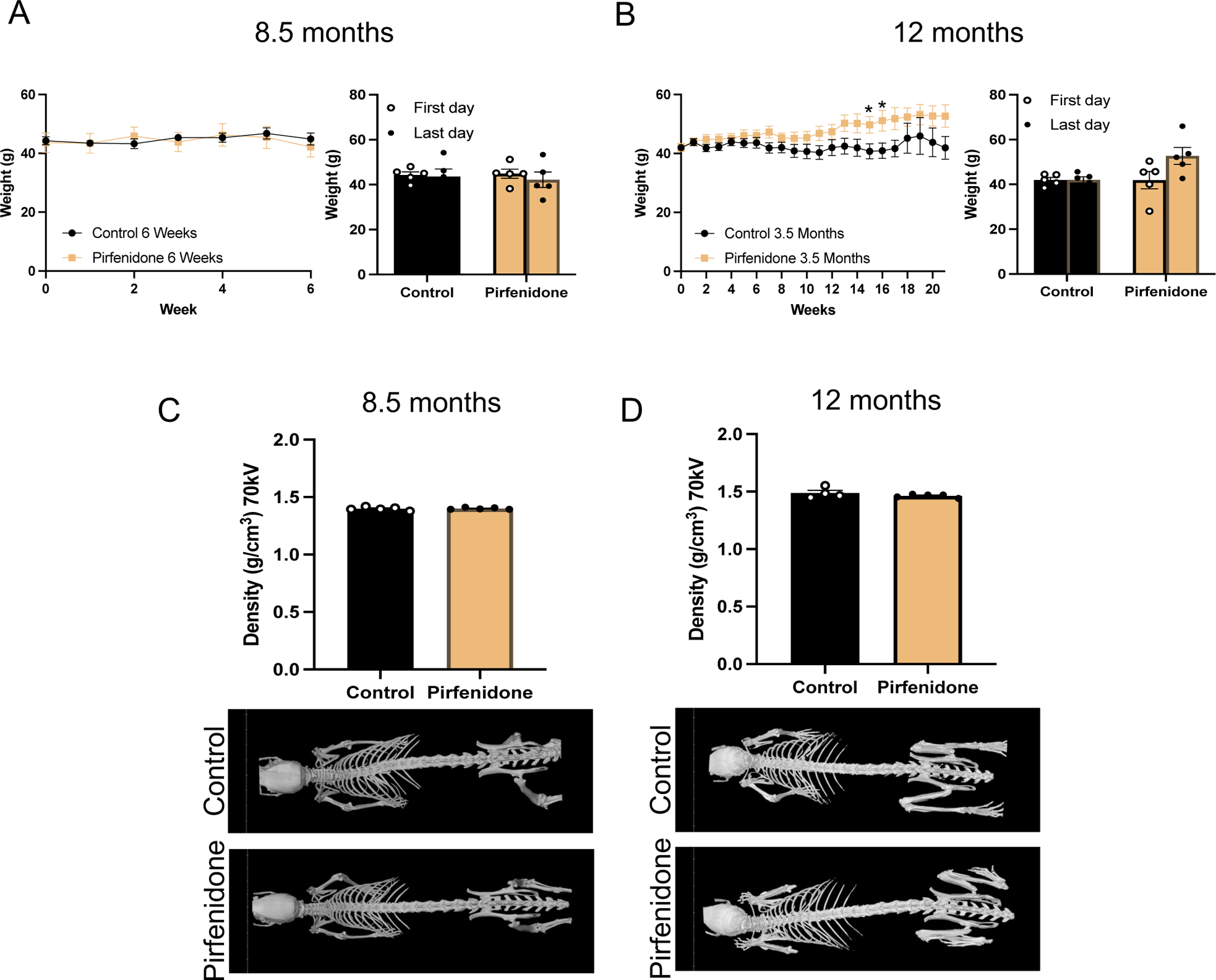
Pirfenidone-treated mice do not show evidence of *in vivo* toxicity based on weight and bone mass density. (A) The graph on the left shows weekly animal weight over 6 weeks of treatment (control or Pirfenidone, 8.5 months). The graph on the right shows the comparison of the average weight of the control and Pirfenidone-treated mice on the first and last days that the measurements were taken. (B) The graph on the left shows weekly animal weight over the time course of 5 months (control or Pirfenidone, 12 months). The graph on the right shows the comparison of the average weight of the control and Pirfenidone-treated mice on the first and last days that the measurements were taken. (C-D) Representative images of bone mass density scans and graphs showing the density quantification in control and Pirfenidone-treated mice (C) immediately after the 6 week treatment (8.5 months) or (D) 3.5 months after the treatment (12 months). Significant differences are noted (** indicates p<0.01).

Given that osteoporosis, or the loss of BMD, is a common age-related phenomenon [58], we wanted to assess whether Pirfenidone affected BMD. Therefore, we performed microCT scans on control and Pirfenidone-treated mice and calculated whole bone density (Figure 4C-D). Both cohorts of mice at 8.5 and 12 months of age had similar BMD measurements (Figure 4C-D). Together our results demonstrate that low dose Pirfenidone exposure can modulate ovarian fibrosis without impacting broad systemic parameters.

### Low dose Pirfenidone treatment improves ovarian morphology and endocrine function

To determine whether the reduction in age-associated fibrosis influences ovarian endpoints, we first examined ovarian histology. Ovaries from control and Pirfenidone treated mice exhibited similar morphology at 8.5 months with follicles and CL present in the tissue (Figure 5A). However, when we quantified the number of follicles, there were more present in the ovaries from the Pirfenidone treated mice relative to controls (p≤0.05, Figure 5B). By 12 months, the number of follicles was similar between the two cohorts. However, the ovarian architecture of Pirfenidone treated mice appeared healthier, with a normal appearing stroma consistent with the decreased accumulation of collagen (Figure 5A). Interestingly, there was a tendency towards a higher number of CLs in ovaries from Pirfenidone-treated mice suggesting improved ovulation (Figure 5B).

**Figure 5:**
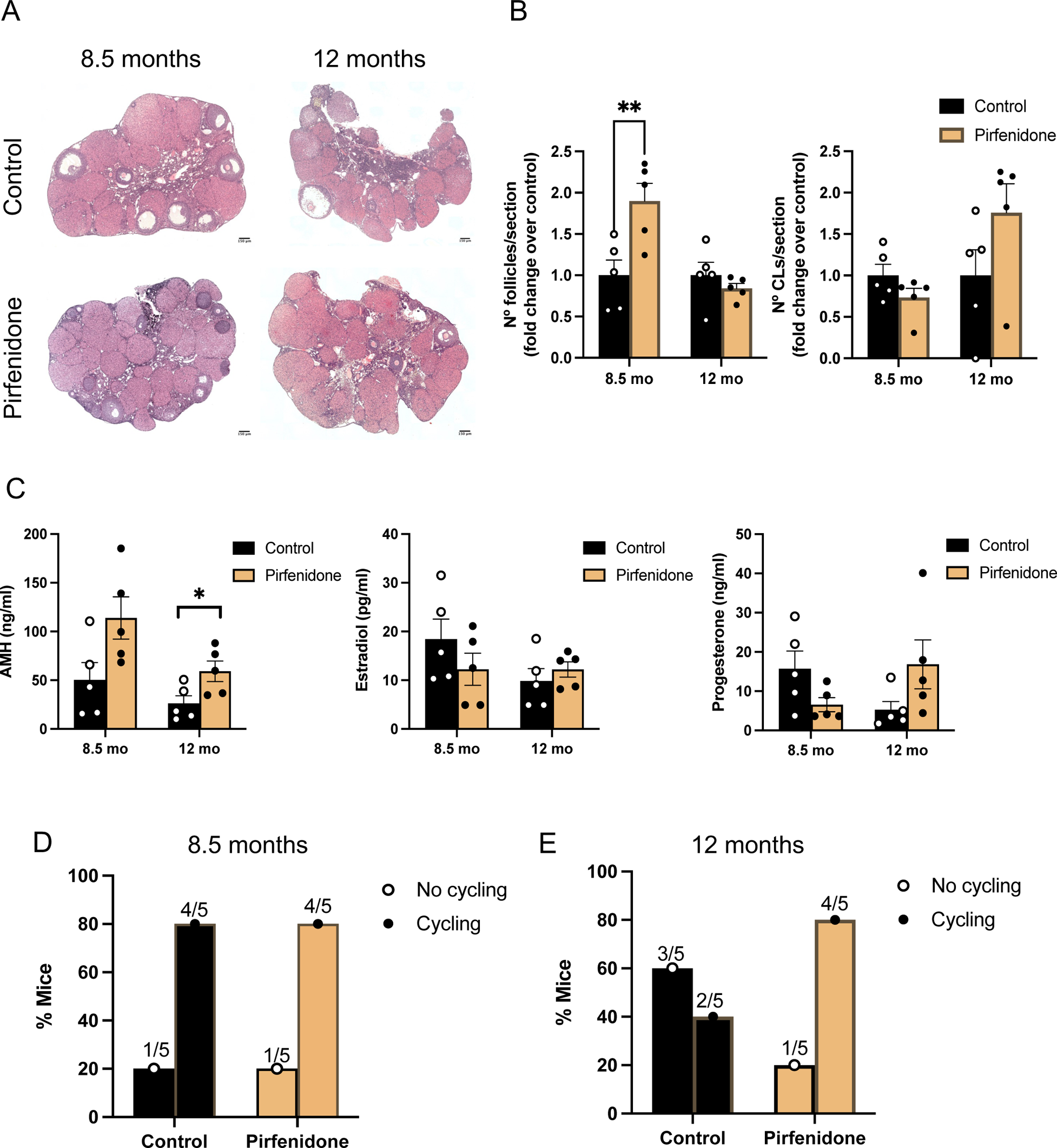
Multiple parameters of ovarian function are significantly improved in Pirfenidone-treated mice. (A) Representative H&E-stained histological sections of ovaries from control and Pirfenidone-treated mice at 8.5 months and 12 months of age. (B) Graphs represent the fold change of the number of follicles and corpora lutea (CL) per ovarian section over control mice at 8.5 months and 12 months of age. (C) Quantification of serum levels of AMH, estradiol, and progesterone in control and Pirfenidone-treated mice at 8.5 months and 12 months. (D-E) Graphs of the percentage of mice that completed a whole estrous cycle during a 15 day period (cycling) compared to those that did not (no cycling) at 8.5 months (D) and 12 months (E). Significant differences are noted (* indicates p<0.05 and ** indicates p<0.01). The scale bar is 150 μm.

To determine whether these histological findings correlated with improved outputs of ovarian function, we assessed circulating gonadal hormone levels. AMH and estradiol are produced by growing follicles, whereas progesterone is produced by CLs and is indicative of ovulation [2, 22]. AMH levels were higher in Pirfenidone-treated mice at both 8.5 and 12 months relative to controls, which is consistent with the larger number of follicles present at 8.5 months (p≤0.05 (only for 12 months mice), Figure 5C). Although there were no differences in estradiol across the cohorts, we observed a clear trend of increased progesterone at 12 months in the Pirfenidone-treated mice compared to controls which is consistent with the increased number of CLs observed in the tissue (Figure 5C).

To determine whether these hormone levels translated into differences in ovarian function, we assessed estrous cyclicity. As expected in reproductively adult females, we observed that 80% the control and Pirfenidone-treated mice at 8.5 months cycled normally (Figure 5D). However, by 12 months only 40% of mice in the control group exhibited normal cyclicity as expected given their advanced age. In contrast, this reduction was not observed in the Pirfenidone-treated mice, where 80% of the mice still exhibited normal cycles (Figure 5E). Taken together, these results suggest that Pirfenidone treatment improves multiple parameters of ovarian function relative to control mice.

### Pirfenidone treatment preserves the ovarian transcriptome and reduces ovarian inflammaging signatures

To further probe the mechanisms that underlie the Pirfenidone-mediated improvements in ovarian function, we performed bulk RNAseq on ovaries to compare gene expression differences between Pirfenidone-treated and control mice at 8.5- and 12-months of age. Volcano plots show that at 8.5 months, 803 and 704 genes were upregulated and downregulated, respectively, in ovaries from Pirfenidone-treated mice compared to controls (Figure 6A). We further examined these genes by performing GO and KEGG analysis (Figure 6B). Downregulated genes were enriched in categories largely related to autophagy, oxidative phosphorylation, and metabolic pathways (Figure 6B), which are enriched in mouse ovaries of advanced reproductive age in other studies [59–61]. Interestingly, genes enriched in pathways related to mitosis and meiosis, cell cycle, ribosome biogenesis, female gamete generation, and regulation of reproductive processes were upregulated which are all indicative of active organ function. From those genes, we identified 9 that are expressed specifically in the female reproductive system and are well-known regulators of folliculogenesis, oogenesis and fertilization (Table 1). These genes include *Amh*, *Gdf9,* and *Zp3*. This data is consistent with the observation that AMH levels were higher in the serum of Pirfenidone-treated mice at 8.5 months, the ovaries had a healthier morphology and a higher number of follicles (Figure 5 A-C).

**Figure 6:**
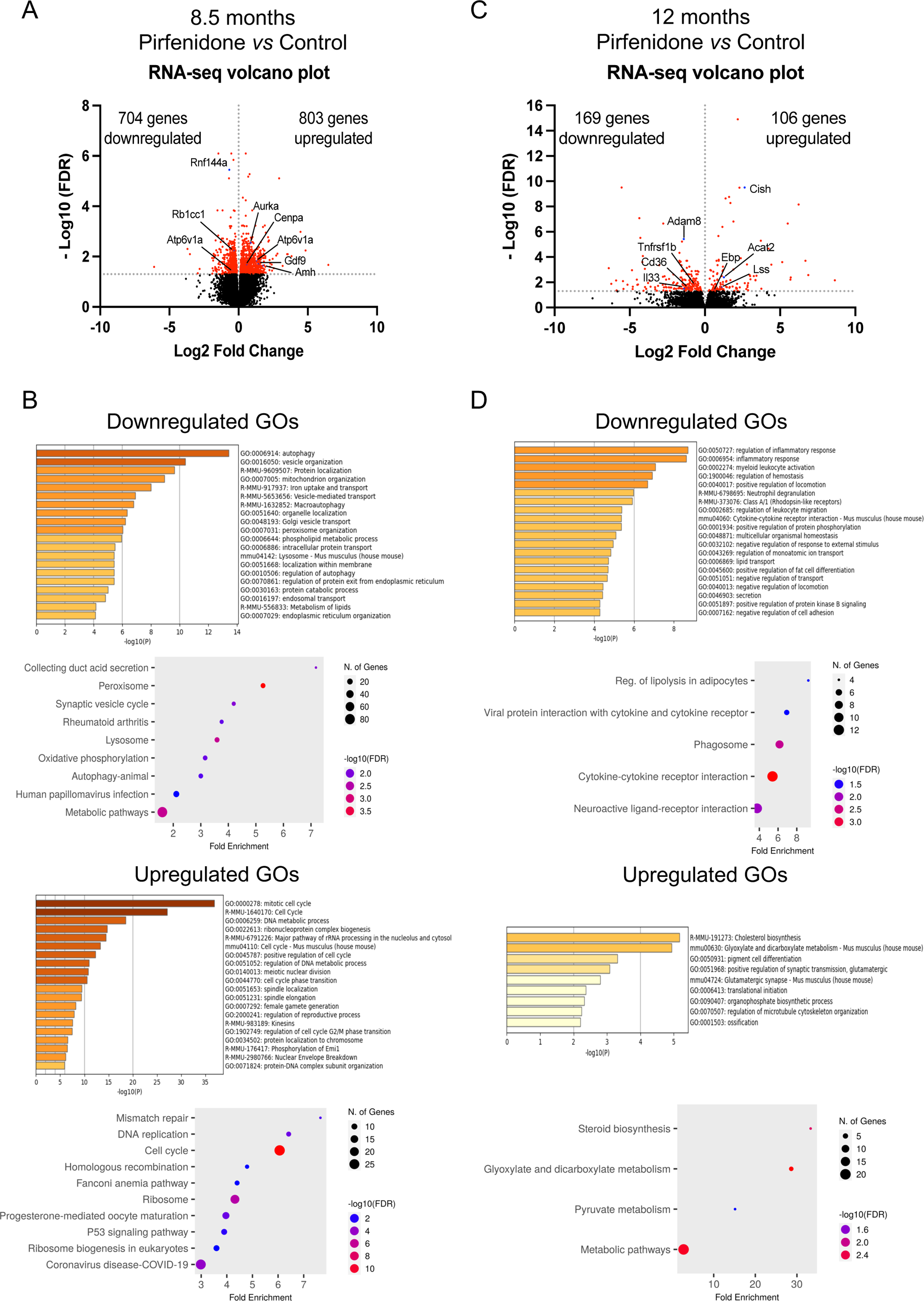
Genes involved in age-associated reduction in ovarian function are downregulated in Pirfenidone-treated mice. (A) A volcano plot of the DEGs between control and Pirfenidone-treated mice at 8.5 months. Of the total 1507 genes that were differently expressed between treatments, 704 were downregulated and 803 upregulated in the Pirfenidone group compared to control. (B, C) Gene Ontology enrichment (bar plot) and KEGG pathway analysis (dot plot) of downregulated (B) and upregulated (C) genes in ovaries from Pirfenidone-treated mice compared to controls at 8.5 months. (D) A volcano plot of the DEGs between control and Pirfenidone-treated mice at 12 months. Of the total 275 genes that were differently expressed between treatments, 169 genes were downregulated and 106 genes were upregulated. (E, F) Gene Ontology (GO) enrichment (bar plot) and KEGG pathway (dot plot) analysis of downregulated (E) and upregulated (F) genes in ovaries from Pirfenidone-treated mice compared to controls at 12 months.

**Table 1:** Upregulated genes specifically involved in female reproductive function in Pirfenidone-treated mice at 8.5 months.

| **Gene** | **Gene description** | **Role** | **Log<sub>2</sub> Fold Change** | **p-value** | **Reference** |
| --- | --- | --- | --- | --- | --- |
| *Amh* | Anti-Mullerian hormone | Folliculogenesis | 1.73 | 0.023 | [1] |
| *Gdf9* | Growth Differentiation Factor 9 | Folliculogenesis | 1.42 | 0.02 | [2, 3] |
| *Nobox* | NOBOX oogenesis Homeobox | Oogenesis | 1.35 | 0.03 | [4, 5] |
| *Zp3* | Zona Pellucida Glycoprotein 3 | Fertilization | 1.33 | 0.01 | [6] |
| *Sgo2a* | Shugoshin 2 | Meiosis | 0.91 | 0.03 | [7] |
| *Zar1* | Zygote Arrest 1 | Maternal factor. Embryogenesis | 1.57 | 0.008 | [8, 9] |
| *Kit* | KIT Proto-Oncogene, Receptor Tyrosine Kinase | Oogenesis and folliculogenesis | 1.28 | 0.04 | [10] |
| *Sycp1* | Synaptonemal Complex Protein 1 | Meiosis | 6.49 | 0.02 | [11] |
| *Izumo1r* | IZUMO1 Receptor, JUNO | Fertilization | 1.49 | 0.01 | [12, 13] |

Remarkably, differentially expressed genes were still present when comparing ovaries from Pirfenidone-treated and control mice at 3.5 months post-treatment (106 genes were upregulated and 169 genes were downregulated in Pirfenidone vs control mice, respectively; Figure 6C). There was reduced expression of genes related to immune and inflammatory pathways (GO: regulation of inflammatory response and inflammatory response; KEGG: viral protein interaction with cytokine and cytokine receptor and cytokine-cytokine receptor interaction). In fact, when we analyzed the genes present in the GO terms related to inflammation, we identified 10 genes whose expression is associated with senescence, accelerated aging, or aging-associated pathologies, and their depletion or decreased expression is correlated with tissue rejuvenation (Table 2). This decrease in inflammation mediated by Pirfenidone appeared to be ovary-specific as we did not observe differences in serum cytokine levels (Supplementary Figure 3). In addition to a decrease in inflammation, ovaries from Pirfenidone treated mice exhibited a higher expression of hormone-related genes relative to controls (GO: cholesterol biosynthesis; KEGG: steroid biosynthesis) (Figure 6D). Altogether, these analyses suggest that Pirfenidone treatment maintains gene expression signatures relevant to ovarian function, including folliculogenesis, oocyte maturation, and ovarian hormone production, while decreasing those associated with ovarian aging, in particular those related to inflammation. Furthermore, Pirfenidone-treatment preserved the ovarian transcriptome across age as there was a 3-fold increase in DEGs in control mice between 8.5 and 12 months compared to Pirfenidone-treated mice (Supplemental Figure 4). Thus, the overall reduction in age-associated transcriptional changes combined with the decreased inflammaging signature observed in ovaries from Pirfenidone-treated mice compared to controls likely contributes to the observed improved ovarian function outcomes.

**Table 2:** Downregulated genes at 12 months in the Pirfenidone-treated mice with well-known age-associated roles in other tissues.

| Gene | Gene description | Role | Log<sub>2</sub> Fold Change | p-value | Reference |
| --- | --- | --- | --- | --- | --- |
| *Adam8* | ADAM Metallopeptidase Domain 8 | Upregulated across many aging tissues, including liver, and species | -1.41 | 3.75E-06 | [14] |
| *Fabp4* | Fatty Acid Binding Protein 4 | Liver rejuvenation when FABP4 is depleted | -1.56 | 0.001 | [15] |
| *Ctss* | Cathepsin S | Involved in age-mediated dry eye | -1.03 | 0.04 | [16] |
| *Fcer1a* | FC Epsilon Receptor 1a | Declined expression in aged human testis and associated with macrophage senescence | -6.40 | 0.0008 | [17] |
| *Serpine1* | Serpin Family E Member 1 | Mutations in *SERPINE1* gene protects against aging in humans | -1.71 | 4.72E-05 | [18] |
| *Cd36* | CD36 Molecule | Expression of CD36 is involved in the mouse age-induced cardiomyopathy | -1.12 | 0.03 | [19] |
| *Cyba* | Cytochrome B-245 Alpha Chain | Age-associated increase expression in multiple tissues and species. Involved in oxidative stress response and senescence | -0.699 | 0.008 | [20] |
| *Ly6d* | Lymphocyte Antigen 6 Family Member D | Involved in mechanisms of senescence cells survival | -4.10 | 0.04 | [21] |
| *Slc7a11* | Solute Carrier Family 7 Member 11 | Inhibition of Slc7a11 ameliorates the profibrotic and senescence phenotype in old lung fibroblasts | -1.50 | 0.01 | [22] |
| *Gdf15* | Growth Differentiation Factor 11 | Higher expression levels in aged humans (human aging biomarker) | -1.58 | 0.0005 | [23] |

## Discussion

Fibrosis and inflammation are hallmarks of many aging tissues, and this is also true of the ovary which ages earlier relative to most organs [5, 10, 11, 15, 20–22, 24–27]. In this study, we advanced the field by demonstrating that we could mitigate age-related ovarian fibrosis through a systemic and constant administration of low dose Pirfenidone, an anti-fibrotic drug, for 6 weeks beginning in mice at 7 months of age when fibrotic foci are first observed in the ovary [20]. With this treatment, we extended ovarian function, since mice treated with Pirfenidone had a higher total number of follicles and CLs, increased ovarian hormone levels, and sustained estrous cyclicity at 12 months of age compared to controls. Interestingly we detected higher levels of ovarian AMH transcripts, serum AMH levels, and total number of growing follicles in Pirfenidone-treated mice compared to controls, suggesting that reducing the fibroinflammatory ovarian microenvironment may provide a more permissive milieu for follicle development. In fact, reducing fibrosis is likely to change the biomechanical properties of the ovary, and follicles grown in less stiff environments have improved follicle function and gamete quality outcomes [29]. Studies are ongoing to determine whether Pirfenidone treatment reduces ovarian stiffness and improves fertility.

Pirfenidone is an FDA-approved broad spectrum anti-fibrotic drug used to treat idiopathic pulmonary fibrosis, and it also inhibits the progression of liver and cardiac fibrosis in various animal models [51–53]. Although Pirfenidone’s mechanism of action is still not completely understood, it exhibits anti-fibrotic properties by decreasing TGF-β1 synthesis and secretion [51–53, 62]. Interestingly, our transcriptomic analysis demonstrated that only 6 weeks of Pirfenidone treatment elicited an enrichment of gene pathways related to oocyte maturation and molecular processes that can protect against age-related oocyte quality decline, including mismatch repair and meiotic regulation. This improvement may be due to reduced reactive oxygen species (ROS) generation, which is a consequence of Pirfenidone treatment in induced lung injury [63–65]. In the ovary, it is well-documented that age-dependent ROS production induces granulosa apoptosis, oocyte death, and reduced reproductive outcomes. ROS are produced in the mitochondria during oxidative phosphorylation and we detected reduced expression of OXPHOS-associated genes in Pirfenidone-treated mice at 8.5 months of age [66]. At 12 months of age, we detected attenuated ovarian inflammatory gene expression in Pirfenidone-treated mice compared to controls. Notably, there was a decrease in genes previously described as overexpressed in the aging ovary, including several TNF receptors, and *Ccl6* (chemokine ligand 6). Increased ovarian inflammaging, and, in particular, enhanced TNFα signalling, is associated with decreased follicle quantity and quality [17]. For example, TNFα promotes follicle death, whereas TNFα knockout mice exhibit increased follicle number, ovulation rate, and overall fertility [67, 68]. Our results are consistent with these findings in that we also observe a clear association between attenuation of inflammaging and enhanced ovarian function, including upregulated expression of genes related to ovarian steroid synthesis and increased hormone production, ovulation capacity, and estrous cyclicity in mice treated with Pirfenidone at 12 months of age.

Our work provides important preclinical evidence that drugs with anti-fibrotic properties could be beneficial in the context of ovarian aging. In fact, our findings are highly complementary to previously published work demonstrating that acute exposure to Pirfenidone can reduce ovarian fibrosis and partially restore the age-dependent defect in ovulation [22, 28]. However, as we move this technology towards clinical translation, there are several important considerations. Strategies that aim to extend ovarian function by reversing the aging process will be complex. Instead, preventative approaches whereby treatment is initiated when ovarian fibrosis and inflammation are first observed, are likely to be more effective. The ovary is one of the first organs to age which enables use of a systemic approach without broadly affecting other tissues. In fact, our serum analysis showed that only ovarian inflammatory markers were impacted after Pirfenidone therapy. Lower doses of drugs could also, in theory, be used over a longer period of time, reducing chances of off target or side effects. Another important consideration is safety. Although Pirfenidone is effective in the rodent, it has significant side effects in humans, including liver damage, which would preclude its use for ovarian indications [57]. In our study, Pirfenidone was administered at very low doses and it did not induce mouse liver toxicity. However, safer alternatives might be needed, and promising preclinical data exist demonstrating that metformin can reduce ovarian fibrosis, alter gene expression in stromal fibroblasts, and improve ovarian function [12, 28]. Thus, further studies validating use of metformin for ovarian longevity are warranted and ongoing. In addition, *ex vivo* models including culture of primary ovarian stromal cells and ovarian explants will enable screening of compounds and expand identification of novel pathways that regulate ovarian fibrosis.

Targeting ovarian fibrosis and stiffness therapeutically has implications beyond reproductive aging as several other conditions are also characterized by increased ovarian fibrosis. These conditions include, but are not limited, to polycystic ovarian syndrome, premature ovarian insufficiency, late effects of chemotherapy and radiation, and endometrioma [69–74]. Moreover, ultrasound-based methods such shear wave elastography can quantify human ovarian stiffness non-invasively which will be essential to evaluate the effectiveness of emerging drugs targeting ovarian fibrosis [75].

---

**Supplementary Figure 1:**
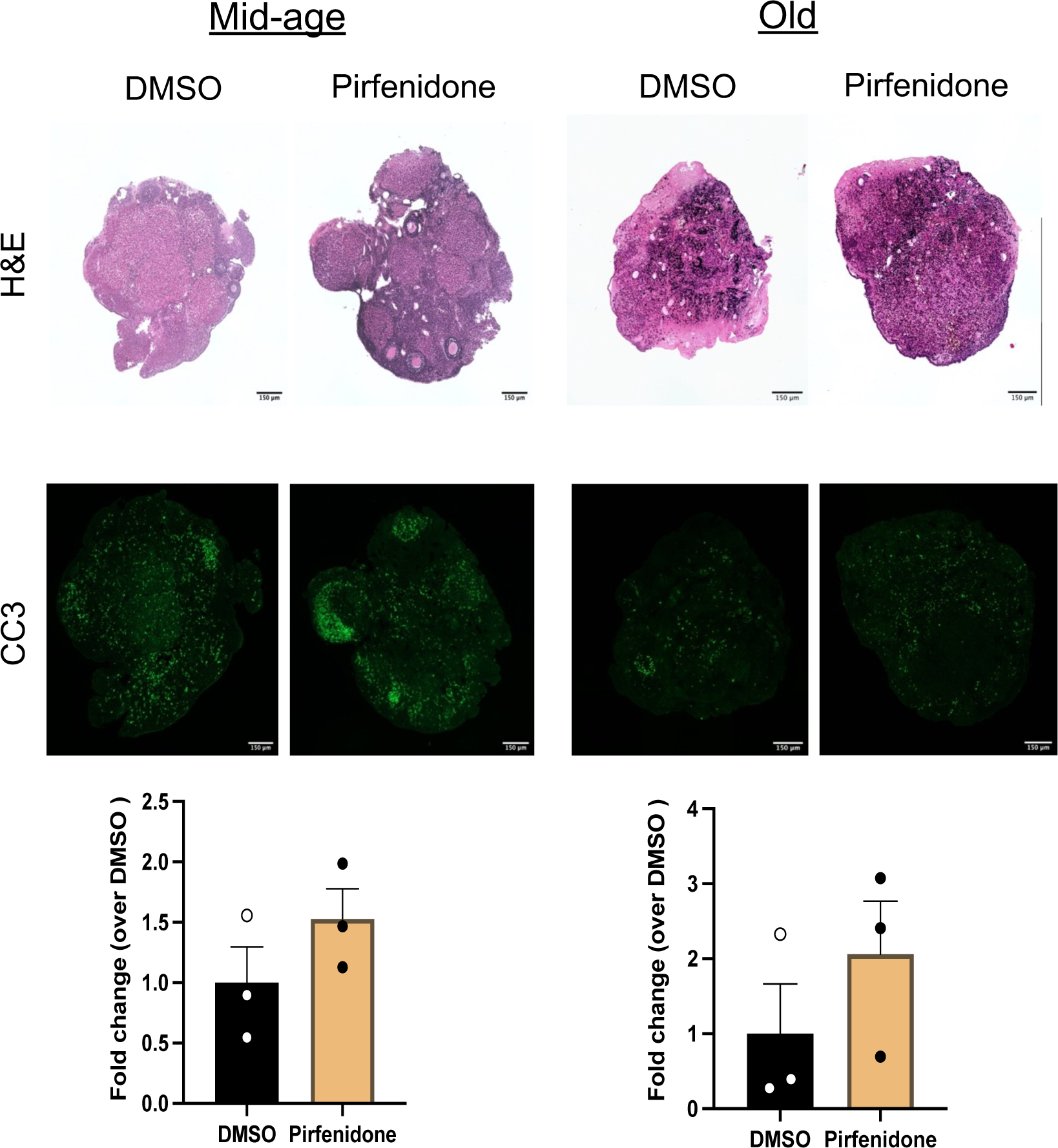
Etanercept and Pirfenidone do not exhibit overt ovarian toxicity *in vitro*. Histological sections of ovarian tissue explants from reproductively mid-age and old mice treated with H_2_O (diluent) or Etanercept (drug) were stained with H&E, and Cleaved Caspase 3 (CC3). The graphs on the bottom show the CC3 intensity in histological sections from ovarian explants from reproductively mid-age and old mice. The scale bar is 150 μm.

**Supplementary Figure 2:**
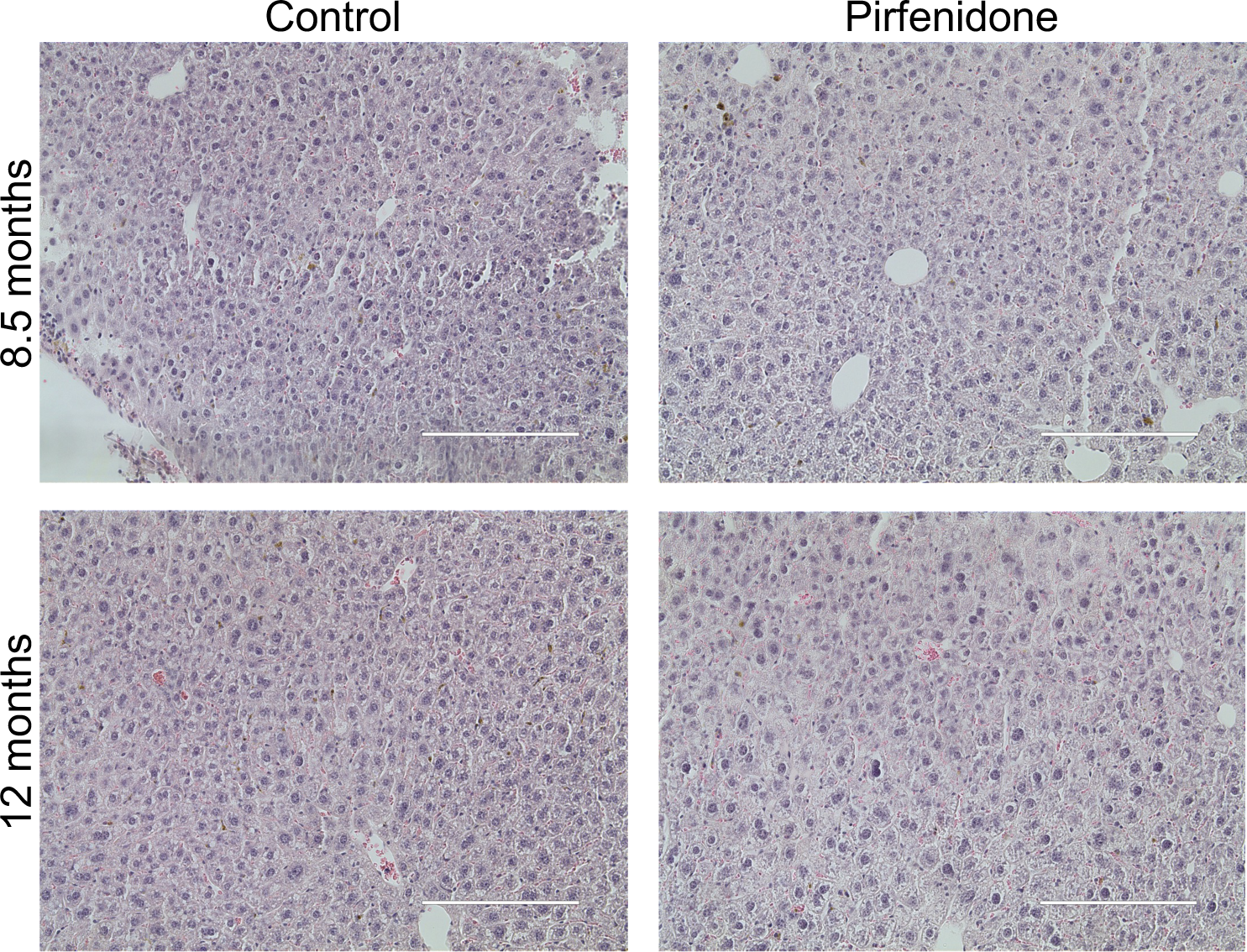
Livers from Pirfenidone-treated mice do not show signs of toxicity. Histological sections of liver tissues from control and Pirfenidone-treated mice at 8.5 months and 12 months stained with H&E. Across the liver lobule from the central vein (CV) to portal triad (PT), there is no evidence of steatosis, necrosis, or inflammatory infiltrate. The scale bar is 200 μm.

**Supplementary Figure 3:**
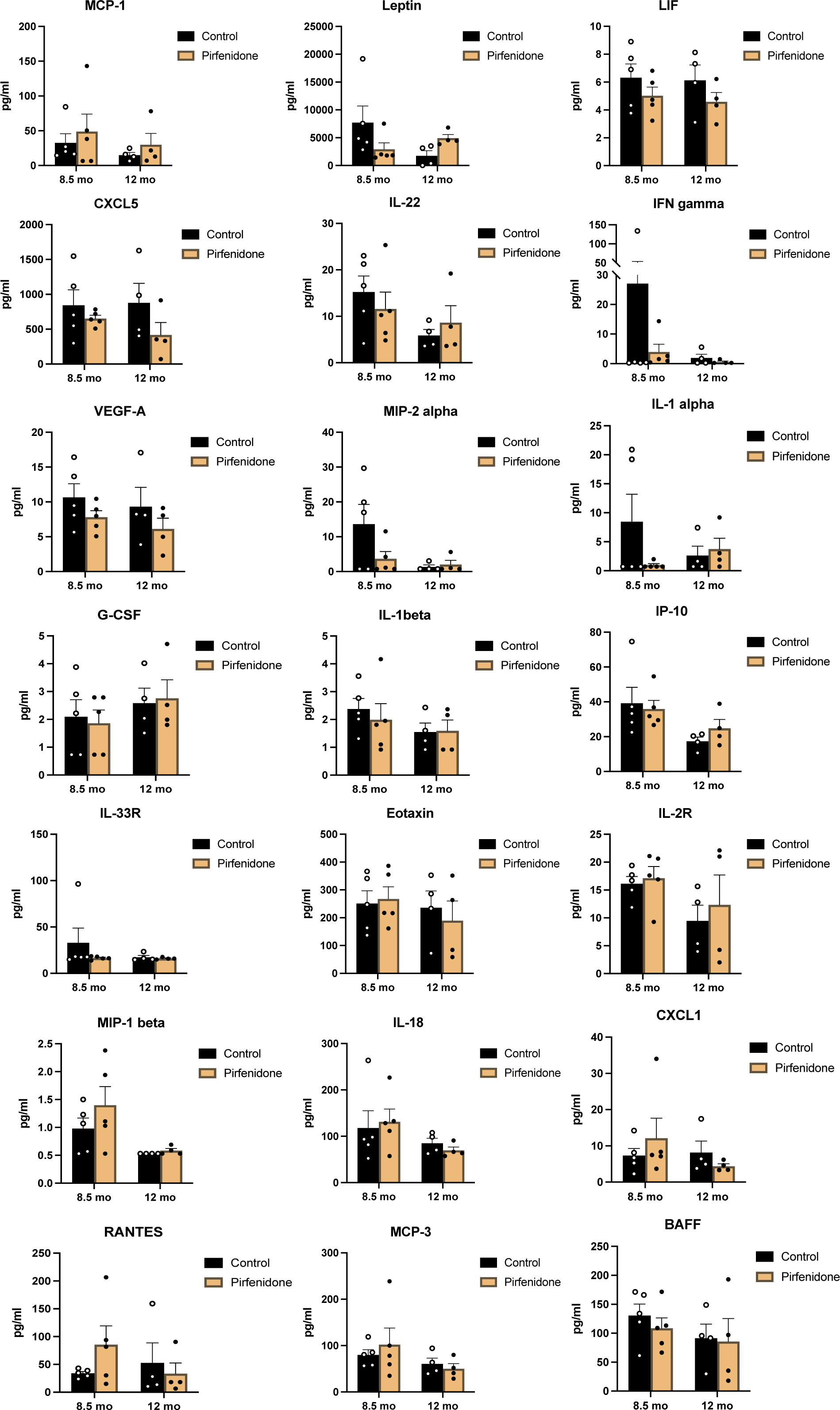
Cytokines, chemokines, and growth factor protein levels in the serum do not change upon Pirfenidone treatment. Graphs are shown that represent the concentration of 21 cytokines, chemokines, and growth factors in the serum of both control and Pirfenidone-treated mice at 8.5 and 12 months.

**Supplementary Figure 4:**
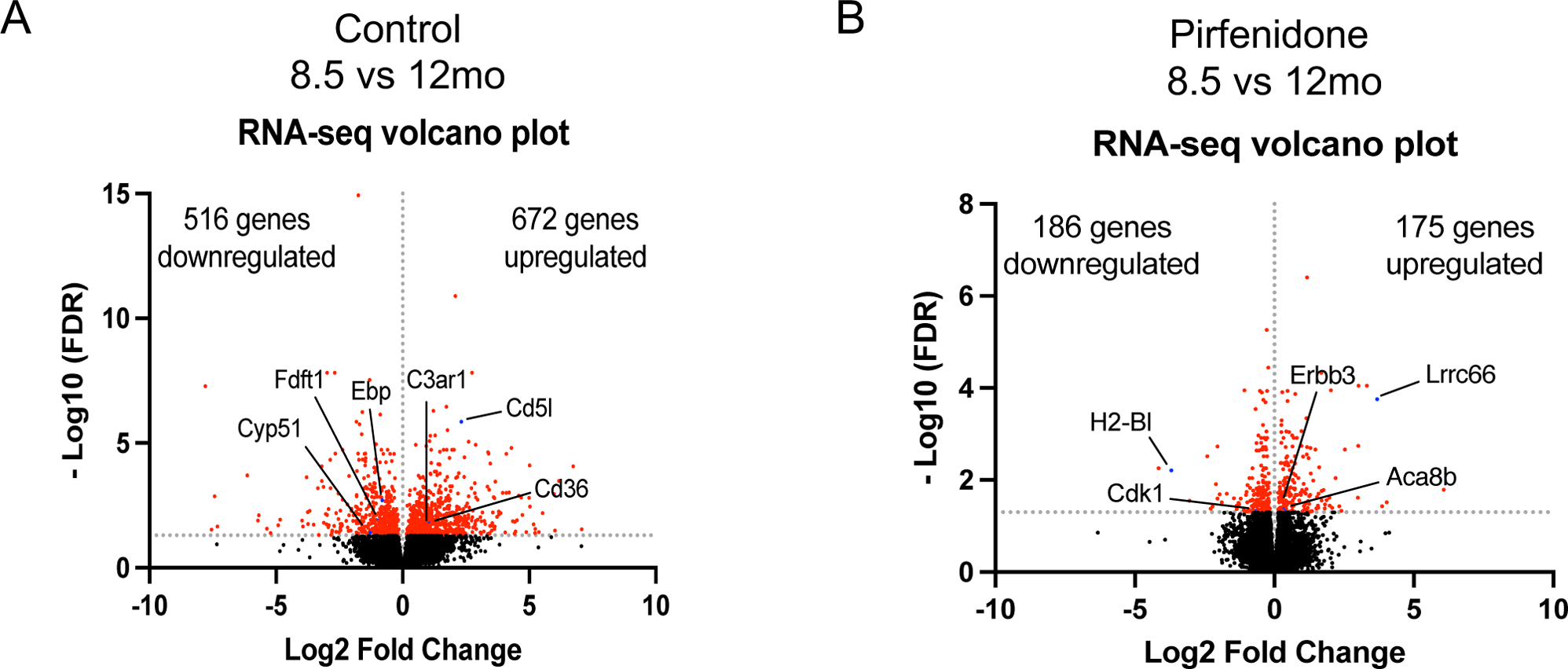
Pirfenidone treatment preserves the ovarian transcriptome from age-associated changes. (A) A volcano plot is shown of the DEGs in control mice at 8.5 and 12 months. Of the total 1188 genes that were differently expressed between ages, 516 were downregulated and 672 upregulated. (B) A volcano plot is shown of the DEGs between Pirfenidone-treated mice at 8.5 and 12 months. Of the total 361 genes that were differently expressed between timepoints, 186 genes were downregulated and 175 genes were upregulated.

**Supplementary Table 1:**
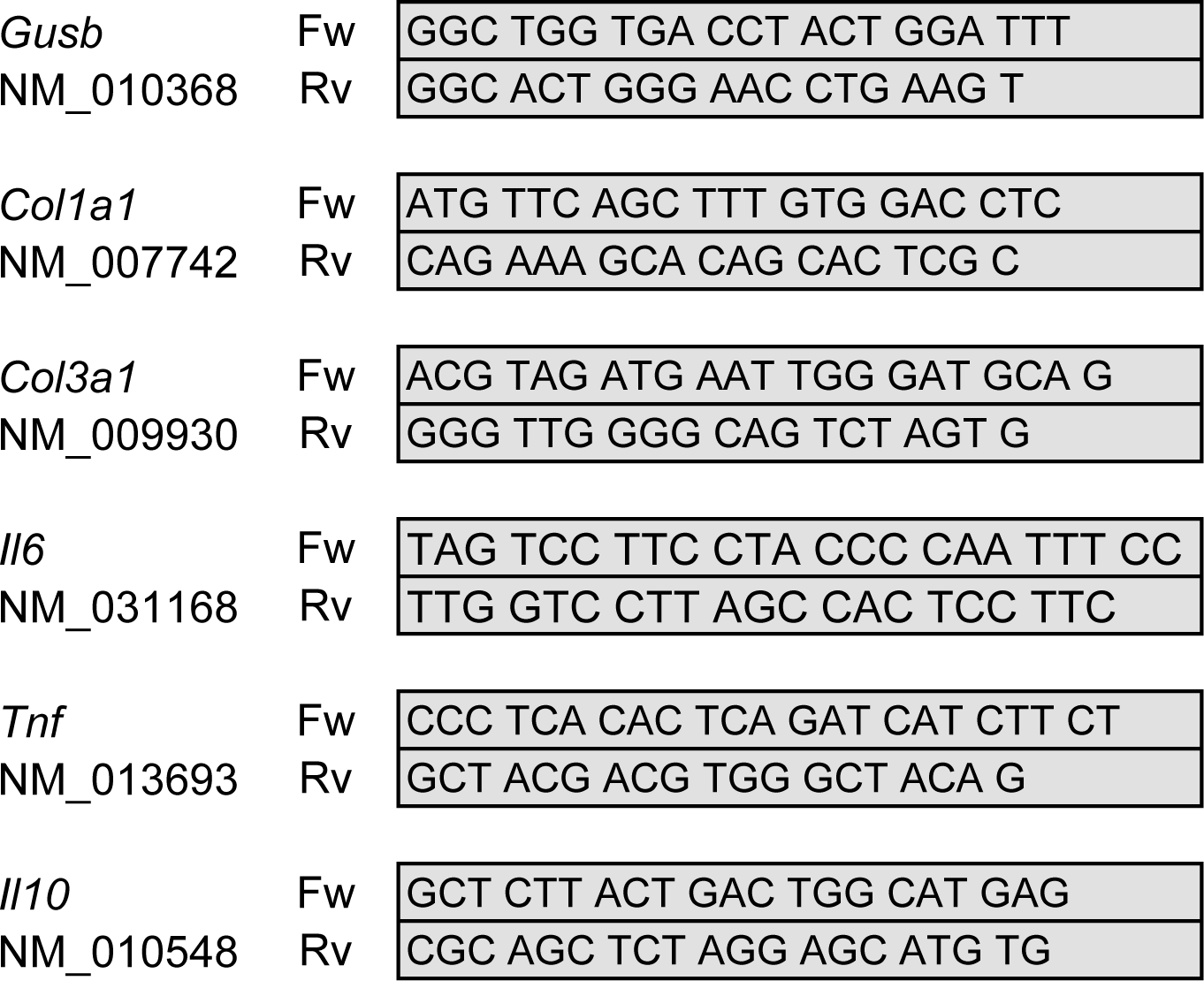
Primers used for RT-qPCR.

## Author contributions

F.A and F.E.D conceived the study, designed the methodology, were involved in funding acquisition, and wrote the manuscript. F.A and C.V performed experiments, data analysis, and visualization. M.T.P. Provided expert feedback, reviewed and edited the manuscript. All authors have read and agreed to the published version of the manuscript.

## Data availability statement

All data generated in this study are available upon request from the corresponding authors. RNA-seq will be available on Gene Expression Omnibus upon publication.

## Acknowledgements

We would like to acknowledge all members of the Duncan laboratory for their support and insightful discussion regarding this work. We thank Karen Velez for her help monitoring the mice. We would like to thank Dr. Hernan E. Lara from the Universidad de Chile for his advice on implanting the osmotic pump. Surgical services were provided by Jiao-Jing Wang from the Northwestern University Comprehensive Transplant Center Microsurgery Core. The RNAseq analysis was supported by the Northwestern University NUSeq Core Facility. The microCT analysis was performed at the CAMI facility, and the multiplex assay was supported by the Comprehensive Metabolic Core at Northwestern University. The ovarian hormone analysis was performed by the Ligand Assay & Analysis Core at the Center for Research in Reproduction from the University of Virginia. Finally, we also thank Dr. John Varga for his critical comments on this work.

## Abbreviations

ECM: Extracellular Matrix
RT: Room Temperature
LPS: Lipopolysaccharide
FBS: Fetal Bovine Serum
FSH: Follicle Stimulating Hormone
PBS: Phosphate Buffered Saline
O/N: Over Night
H&E: Haematoxylin & Eosin
PSR: Picrosirius Red
TBS: Tris Buffered Saline
ABC: Avidin-Biotin Complex
BMD: Bone Mass Density
CL: Corpora Lutea
E2: Estradiol
P4: Progesterone
AMH: anti-Müllerian hormone
FDR: False Discovery Rate
GO: Gene Ontology
ROS: Reactive Oxygen Species.

## References

1. Hainaut, M. and H.J. Clarke, Germ cells of the mammalian female: A limited or renewable resource?dagger. Biol Reprod, 2021. 105(4): p. 774–788.

2. Hsueh, A.J., et al., Intraovarian control of early folliculogenesis. Endocr Rev, 2015. 36(1): p. 1–24.

3. Doherty, C.A., et al., Bidirectional communication in oogenesis: a dynamic conversation in mice and Drosophila. Trends Cell Biol, 2022. 32(4): p. 311–323.

4. Broekmans, F.J., M.R. Soules, and B.C. Fauser, Ovarian aging: mechanisms and clinical consequences. Endocr Rev, 2009. 30(5): p. 465–93.

5. Wu, J., et al., Aging conundrum: A perspective for ovarian aging. Front Endocrinol (Lausanne), 2022. 13: p. 952471.

6. Hagg, S. and J. Jylhava, Sex differences in biological aging with a focus on human studies. Elife, 2021. 10.

7. The Lancet Diabetes, E., Menopause: a turning point for women’s health. Lancet Diabetes Endocrinol, 2022. 10(6): p. 373.

8. Francesca E. Duncan, R.C., Mary Ellen Pavone, Chapter 9 - Female Reproductive Aging: From Consequences to Mechanisms, Markers, and Treatments, in Conn’s Handbook of Models for Human Aging (Second Edition). 2018.

9. Kinnear, H.M., et al., The ovarian stroma as a new frontier. Reproduction, 2020. 160(3): p. R25–R39.

10. Isola, J.V.V., et al., A single-cell atlas of the aging mouse ovary. Nat Aging, 2024. 4(1): p. 145–162.

11. Ben Yaakov, T., et al., Single-cell analysis of the aged ovarian immune system reveals a shift towards adaptive immunity and attenuated cell function. Elife, 2023. 12.

12. Landry, D.A., et al., Metformin prevents age-associated ovarian fibrosis by modulating the immune landscape in female mice. Sci Adv, 2022. 8(35): p. eabq1475.

13. Foley, K.G., M.T. Pritchard, and F.E. Duncan, Macrophage-derived multinucleated giant cells: hallmarks of the aging ovary. Reproduction, 2021. 161(2): p. V5–V9.

14. Zhang, Z., et al., Inflammaging is associated with shifted macrophage ontogeny and polarization in the aging mouse ovary. Reproduction, 2020. 159(3): p. 325–337.

15. Winkler, I., et al., The cycling and aging mouse female reproductive tract at single-cell resolution. Cell, 2024. 187(4): p. 981–998 e25.

16. Lliberos, C., et al., The Inflammasome Contributes to Depletion of the Ovarian Reserve During Aging in Mice. Front Cell Dev Biol, 2020. 8: p. 628473.

17. Lliberos, C., et al., Evaluation of inflammation and follicle depletion during ovarian ageing in mice. Sci Rep, 2021. 11(1): p. 278.

18. Perrone, R., et al., CD38 regulates ovarian function and fecundity via NAD(+) metabolism. iScience, 2023. 26(10): p. 107949.

19. Isola, J.V.V., et al., Inflammation, immune cells, and cellular senescence in the aging ovary. Reproduction, 2024.

20. Briley, S.M., et al., Reproductive age-associated fibrosis in the stroma of the mammalian ovary. Reproduction, 2016. 152(3): p. 245–260.

21. Amargant, F., et al., Ovarian stiffness increases with age in the mammalian ovary and depends on collagen and hyaluronan matrices. Aging Cell, 2020. 19(11): p. e13259.

22. Mara, J.N., et al., Ovulation and ovarian wound healing are impaired with advanced reproductive age. Aging (Albany NY), 2020. 12(10): p. 9686–9713.

23. McCloskey, C.W., et al., Metformin Abrogates Age-Associated Ovarian Fibrosis. Clin Cancer Res, 2020. 26(3): p. 632–642.

24. Ouni, E., et al., A blueprint of the topology and mechanics of the human ovary for next-generation bioengineering and diagnosis. Nat Commun, 2021. 12(1): p. 5603.

25. Machlin, J.H., et al., Fibroinflammatory Signatures Increase with Age in the Human Ovary and Follicular Fluid. Int J Mol Sci, 2021. 22(9).

26. Ouni, E., et al., Proteome-wide and matrisome-specific atlas of the human ovary computes fertility biomarker candidates and open the way for precision oncofertility. Matrix Biol, 2022. 109: p. 91–120.

27. Ouni, E., et al., Spatiotemporal changes in mechanical matrisome components of the human ovary from prepuberty to menopause. Hum Reprod, 2020. 35(6): p. 1391–1410.

28. Umehara, T., et al., Female reproductive life span is extended by targeted removal of fibrotic collagen from the mouse ovary. Sci Adv, 2022. 8(24): p. eabn4564.

29. Pietroforte, S., M. Plough, and F. Amargant, Age-associated increased stiffness of the ovarian microenvironment impairs follicle development and oocyte quality and rapidly alters follicle gene expression. bioRxiv, 2024: p. 2024.06.09.598134.

30. Alkmin, S., M.S. Patankar, and P.J. Campagnola, Assessing the roles of collagen fiber morphology and matrix stiffness on ovarian cancer cell migration dynamics using multiphoton fabricated orthogonal image-based models. Acta Biomater, 2022. 153: p. 342–354.

31. Fujimoto, H., et al., Tumor-associated fibrosis: a unique mechanism promoting ovarian cancer metastasis and peritoneal dissemination. Cancer Metastasis Rev, 2024.

32. Landry, D.A., H.T. Vaishnav, and B.C. Vanderhyden, The significance of ovarian fibrosis. Oncotarget, 2020. 11(47): p. 4366–4370.

33. Dean, M., et al., Exposure of the extracellular matrix and colonization of the ovary in metastasis of fallopian-tube-derived cancer. Carcinogenesis, 2019. 40(1): p. 41–51.

34. Schaefer, C.J., et al., Antifibrotic activities of pirfenidone in animal models. Eur Respir Rev, 2011. 20(120): p. 85–97.

35. Rowley, J.E., et al., Low Molecular Weight Hyaluronan Induces an Inflammatory Response in Ovarian Stromal Cells and Impairs Gamete Development In Vitro. Int J Mol Sci, 2020. 21(3).

36. Parkes, W.S., et al., Hyaluronan and Collagen Are Prominent Extracellular Matrix Components in Bovine and Porcine Ovaries. Genes (Basel), 2021. 12(8).

37. Byers, S.L., et al., Mouse estrous cycle identification tool and images. PLoS One, 2012. 7(4): p. e35538.

38. Dobin, A., et al., STAR: ultrafast universal RNA-seq aligner. Bioinformatics, 2013. 29(1): p. 15–21.

39. Anders, S., P.T. Pyl, and W. Huber, HTSeq--a Python framework to work with high-throughput sequencing data. Bioinformatics, 2015. 31(2): p. 166–9.

40. Love, M.I., W. Huber, and S. Anders, Moderated estimation of fold change and dispersion for RNA-seq data with DESeq2. Genome Biol, 2014. 15(12): p. 550.

41. Tabas-Madrid, D., R. Nogales-Cadenas, and A. Pascual-Montano, GeneCodis3: a non-redundant and modular enrichment analysis tool for functional genomics. Nucleic Acids Res, 2012. 40(Web Server issue): p. W478–83.

42. Nogales-Cadenas, R., et al., GeneCodis: interpreting gene lists through enrichment analysis and integration of diverse biological information. Nucleic Acids Res, 2009. 37(Web Server issue): p. W317–22.

43. Carmona-Saez, P., et al., GENECODIS: a web-based tool for finding significant concurrent annotations in gene lists. Genome Biol, 2007. 8(1): p. R3.

44. Ge, S.X., D. Jung, and R. Yao, ShinyGO: a graphical gene-set enrichment tool for animals and plants. Bioinformatics, 2020. 36(8): p. 2628–2629.

45. Mitoma, H., et al., Molecular mechanisms of action of anti-TNF-alpha agents - Comparison among therapeutic TNF-alpha antagonists. Cytokine, 2018. 101: p. 56–63.

46. Haraoui, B. and V. Bykerk, Etanercept in the treatment of rheumatoid arthritis. Ther Clin Risk Manag, 2007. 3(1): p. 99–105.

47. Tong, W., et al., Resveratrol inhibits LPS-induced inflammation through suppressing the signaling cascades of TLR4-NF-kappaB/MAPKs/IRF3. Exp Ther Med, 2020. 19(3): p. 1824–1834.

48. Honda, T. and H. Inagawa, Utility of In Vitro Cellular Models of Low-Dose Lipopolysaccharide in Elucidating the Mechanisms of Anti-Inflammatory and Wound-Healing-Promoting Effects of Lipopolysaccharide Administration In Vivo. Int J Mol Sci, 2023. 24(18).

49. Thakur, V., et al., Chronic ethanol feeding increases activation of NADPH oxidase by lipopolysaccharide in rat Kupffer cells: role of increased reactive oxygen in LPS-stimulated ERK1/2 activation and TNF-alpha production. J Leukoc Biol, 2006. 79(6): p. 1348–56.

50. Pritchard, M.T., Z. Li, and E.A. Repasky, Nitric oxide production is regulated by fever-range thermal stimulation of murine macrophages. J Leukoc Biol, 2005. 78(3): p. 630–8.

51. Ruwanpura, S.M., B.J. Thomas, and P.G. Bardin, Pirfenidone: Molecular Mechanisms and Potential Clinical Applications in Lung Disease. Am J Respir Cell Mol Biol, 2020. 62(4): p. 413–422.

52. Sartiani, L., et al., Pharmacological basis of the antifibrotic effects of pirfenidone: Mechanistic insights from cardiac in-vitro and in-vivo models. Front Cardiovasc Med, 2022. 9: p. 751499.

53. Nakazato, H., et al., A novel anti-fibrotic agent pirfenidone suppresses tumor necrosis factor-alpha at the translational level. Eur J Pharmacol, 2002. 446(1-3): p. 177–85.

54. Tsukui, T., et al., Collagen-producing lung cell atlas identifies multiple subsets with distinct localization and relevance to fibrosis. Nat Commun, 2020. 11(1): p. 1920.

55. Pappas, L.E. and T.R. Nagy, The translation of age-related body composition findings from rodents to humans. Eur J Clin Nutr, 2019. 73(2): p. 172–178.

56. Benesic, A., K. Jalal, and A.L. Gerbes, Acute Liver Failure During Pirfenidone Treatment Triggered by Co-Medication With Esomeprazole. Hepatology, 2019. 70(5): p. 1869–1871.

57. Risk of serious liver injury with pirfenidone: updated advice. Reactions Weekly, 2020. 1831(1): p. 3–3.

58. Shim, J., et al., Micro-computed tomography assessment of bone structure in aging mice. Sci Rep, 2022. 12(1): p. 8117.

59. Li, F., et al., Targeting Estrogen Receptor Beta Ameliorates Diminished Ovarian Reserve via Suppression of the FOXO3a/Autophagy Pathway. Aging Dis, 2024.

60. Yan, F., et al., The role of oxidative stress in ovarian aging: a review. J Ovarian Res, 2022. 15(1): p. 100.

61. Duncan, F.E., et al., Age-associated dysregulation of protein metabolism in the mammalian oocyte. Aging Cell, 2017. 16(6): p. 1381–1393.

62. Xi, Y., et al., The anti-fibrotic drug pirfenidone inhibits liver fibrosis by targeting the small oxidoreductase glutaredoxin-1. Sci Adv, 2021. 7(36): p. eabg9241.

63. Pourgholamhossein, F., et al., Pirfenidone protects against paraquat-induced lung injury and fibrosis in mice by modulation of inflammation, oxidative stress, and gene expression. Food Chem Toxicol, 2018. 112: p. 39–46.

64. Seifirad, S., et al., Effect of pirfenidone on pulmonary fibrosis due to paraquat poisoning in rats. Clin Toxicol (Phila), 2012. 50(8): p. 754–8.

65. Giri, S.N., et al., Effects of pirfenidone on the generation of reactive oxygen species in vitro. J Environ Pathol Toxicol Oncol, 1999. 18(3): p. 169–77.

66. Bhatti, J.S., G.K. Bhatti, and P.H. Reddy, Mitochondrial dysfunction and oxidative stress in metabolic disorders - A step towards mitochondria based therapeutic strategies. Biochim Biophys Acta Mol Basis Dis, 2017. 1863(5: p. 1066–1077.

67. Morrison, L.J. and J.L. Marcinkiewicz, Tumor necrosis factor alpha enhances oocyte/follicle apoptosis in the neonatal rat ovary. Biol Reprod, 2002. 66(2): p. 450–7.

68. Cui, L.L., et al., Tumor necrosis factor alpha knockout increases fertility of mice. Theriogenology, 2011. 75(5): p. 867–76.

69. Wood, C.D., et al., Multi-modal magnetic resonance elastography for noninvasive assessment of ovarian tissue rigidity in vivo. Acta Biomater, 2015. 13: p. 295–300.

70. Mendez, M., et al., Biomechanical characteristics of the ovarian cortex in POI patients and functional outcomes after drug-free IVA. J Assist Reprod Genet, 2022. 39(8): p. 1759–1767.

71. Erol Koc, E.M., et al., Prolidase as a marker of fibrogenesis in idiopathic primary ovarian insufficiency. Eur J Obstet Gynecol Reprod Biol, 2023. 281: p. 7–11.

72. Amargant, F., et al., Sphingosine-1-phosphate and its mimetic FTY720 do not protect against radiation-induced ovarian fibrosis in the nonhuman primatedagger. Biol Reprod, 2021. 104(5): p. 1058–1070.

73. Shi, L.B., et al., Transforming growth factor beta1 from endometriomas promotes fibrosis in surrounding ovarian tissues via Smad2/3 signaling. Biol Reprod, 2017. 97(6): p. 873–882.

74. Kitajima, M., et al., Endometriomas as a possible cause of reduced ovarian reserve in women with endometriosis. Fertil Steril, 2011. 96(3): p. 685–91.

75. Zaniker, E.J., et al., Shear wave elastography to assess stiffness of the human ovary and other reproductive tissues across the reproductive lifespan in health and diseasedagger. Biol Reprod, 2024.

## References

1. Gruijters, M.J., et al., Anti-Mullerian hormone and its role in ovarian function. Mol Cell Endocrinol, 2003. 211(1-2): p. 85–90.

2. McGrath, S.A., A.F. Esquela, and S.J. Lee, Oocyte-specific expression of growth/differentiation factor-9. Mol Endocrinol, 1995. 9(1): p. 131–6.

3. Dong, J., et al., Growth differentiation factor-9 is required during early ovarian folliculogenesis. Nature, 1996. 383(6600): p. 531–5.

4. Rajkovic, A., et al., NOBOX deficiency disrupts early folliculogenesis and oocyte-specific gene expression. Science, 2004. 305(5687): p. 1157–9.

5. Suzumori, N., et al., Nobox is a homeobox-encoding gene preferentially expressed in primordial and growing oocytes. Mech Dev, 2002. 111(1-2): p. 137–41.

6. Chamberlin, M.E. and J. Dean, Human homolog of the mouse sperm receptor. Proc Natl Acad Sci U S A, 1990. 87(16): p. 6014–8.

7. Kitajima, T.S., S.A. Kawashima, and Y. Watanabe, The conserved kinetochore protein shugoshin protects centromeric cohesion during meiosis. Nature, 2004. 427(6974): p. 510–7.

8. Wu, X., et al., Zygote arrest 1 (Zar1) is an evolutionarily conserved gene expressed in vertebrate ovaries. Biol Reprod, 2003. 69(3): p. 861–7.

9. Wu, X., et al., Zygote arrest 1 (Zar1) is a novel maternal-effect gene critical for the oocyte-to-embryo transition. Nat Genet, 2003. 33(2): p. 187–91.

10. Horie, K., et al., The expression of c-kit protein in human adult and fetal tissues. Hum Reprod, 1993. 8(11): p. 1955–62.

11. Paredes, A., et al., Loss of synaptonemal complex protein-1, a synaptonemal complex protein, contributes to the initiation of follicular assembly in the developing rat ovary. Endocrinology, 2005. 146(12): p. 5267–77.

12. Bianchi, E., et al., Juno is the egg Izumo receptor and is essential for mammalian fertilization. Nature, 2014. 508(7497): p. 483–7.

13. Wang, H., X. Hong, and W.H. Kinsey, Sperm-oocyte signaling: the role of IZUMO1R and CD9 in PTK2B activation and actin remodeling at the sperm binding sitedagger. Biol Reprod, 2021. 104(6): p. 1292–1301.

14. Naus, S., et al., The metalloprotease-disintegrin ADAM8 is essential for the development of experimental asthma. Am J Respir Crit Care Med, 2010. 181(12): p. 1318–28.

15. Lv, J., et al., Targeting FABP4 in elderly mice rejuvenates liver metabolism and ameliorates aging-associated metabolic disorders. Metabolism, 2023. 142: p. 155528.

16. Yu, Z., et al., Cathepsin S is a novel target for age-related dry eye. Exp Eye Res, 2022. 214: p. 108895.

17. Nie, X., et al., Single-cell analysis of human testis aging and correlation with elevated body mass index. Dev Cell, 2022. 57(9): p. 1160–1176 e5.

18. Khan, S.S., et al., A null mutation in SERPINE1 protects against biological aging in humans. Sci Adv, 2017. 3(11): p. eaao1617.

19. Koonen, D.P., et al., CD36 expression contributes to age-induced cardiomyopathy in mice. Circulation, 2007. 116(19): p. 2139–47.

20. Barth, E., et al., Conserved aging-related signatures of senescence and inflammation in different tissues and species. Aging (Albany NY), 2019. 11(19): p. 8556–8572.

21. Nagano, T., et al., LY6D-induced macropinocytosis as a survival mechanism of senescent cells. J Biol Chem, 2021. 296: p. 100049.

22. Ritzenthaler, J.D., et al., The profibrotic and senescence phenotype of old lung fibroblasts is reversed or ameliorated by genetic and pharmacological manipulation of Slc7a11 expression. Am J Physiol Lung Cell Mol Physiol, 2022. 322(3): p. L449–L461.

23. Liu, H., et al., GDF15 as a biomarker of ageing. Exp Gerontol, 2021. 146: p. 111228.

